# Efficient and reproducible pipelines for spike sorting large-scale electrophysiology data

**DOI:** 10.1101/2025.11.12.687966

**Authors:** Alessio P. Buccino, Arjun Sridhar, David Feng, Karel Svoboda, Joshua H. Siegle

## Abstract

The scale of *in vivo* electrophysiology has expanded in recent years, with simultaneous recordings across thousands of electrodes now becoming routine. These advances have enabled a wide range of discoveries, but they also impose substantial computational demands. Spike sorting, the procedure that extracts spikes from extracellular voltage measurements, remains a major bottleneck: a dataset collected in a few hours can take days to spike sort on a single machine, and the field lacks rigorous validation of the many spike sorting algorithms and preprocessing steps that are in use. Advancing the speed and accuracy of spike sorting is essential to fully realize the potential of large-scale electrophysiology. Here, we present an end-to-end spike sorting pipeline that leverages parallelization to scale to large datasets. The same workflow can run reproducibly on individual workstations, high-performance computing clusters, or cloud environments, with computing resources tailored to each processing step to reduce costs and execution times. In addition, we introduce a benchmarking pipeline, also optimized for parallel processing, that enables systematic comparison of multiple sorting pipelines. Using this framework, we show that `Kilosort4`, a widely used spike sorting algorithm, outperforms `Kilosort2.5` (Pachitariu et al. 2024). We also show that 7*×* lossy compression, which substantially reduces the cost of data storage, has minimal impact on spike sorting performance. Together, these pipelines address the urgent need for scalable and transparent spike sorting of electrophysiology data, preparing the field for the coming flood of multi-thousand-channel experiments.

## Introduction

A central goal of systems neuroscience is to connect the spikes of populations of individual neurons to the flow of information across neural circuits and ultimately to behavior (Abbott & Svoboda 2020). Extracellular electrophysiology with implanted electrode arrays is the most widely used method for establishing this link. This approach requires a processing step known as “spike sorting”, in which spikes are separated from noise and assigned to individual neurons (Obien et al. 2015; Harris et al. 2016). Spike sorting is difficult, because spikes last for around a millisecond, their amplitudes attenuate over tens of µm, and they occur in densely packed neural tissue (Einevoll et al. 2012; Gold et al. 2006). Spikes from an individual neuron may be readily distinguished from those of its neighbors within a small radius from the soma, but they become increasingly difficult to identify at greater distances (Henze et al. 2000; Buzsáki 2004). Despite these inherent challenges, accurate spike sorting is essential for uncovering the mechanisms that shape brain-wide patterns of activity.

Scaling up electrophysiology entails adding electrodes with spacing matched to neuron densities (∼20 µm spacing), while maintaining sampling rates high enough to capture the details of spike waveforms (∼30 kHz) (Marblestone et al. 2013; Kleinfeld et al. 2019) (Figure 1a). Thus, the overall size of an electrophysiology dataset increases roughly in proportion to the number of simultaneously recorded neurons. Even with modern hardware acceleration, processing these datasets remains computationally intensive, often taking much longer than the recording itself (Figure 1b). As experiments expand to include more probes and recordings over many days of natural behavior (Campagner et al. 2025; Dhawale et al. 2017; Newman et al. 2025), spike sorting becomes impossible to sustain without large-scale parallelization.

**Figure 1:**
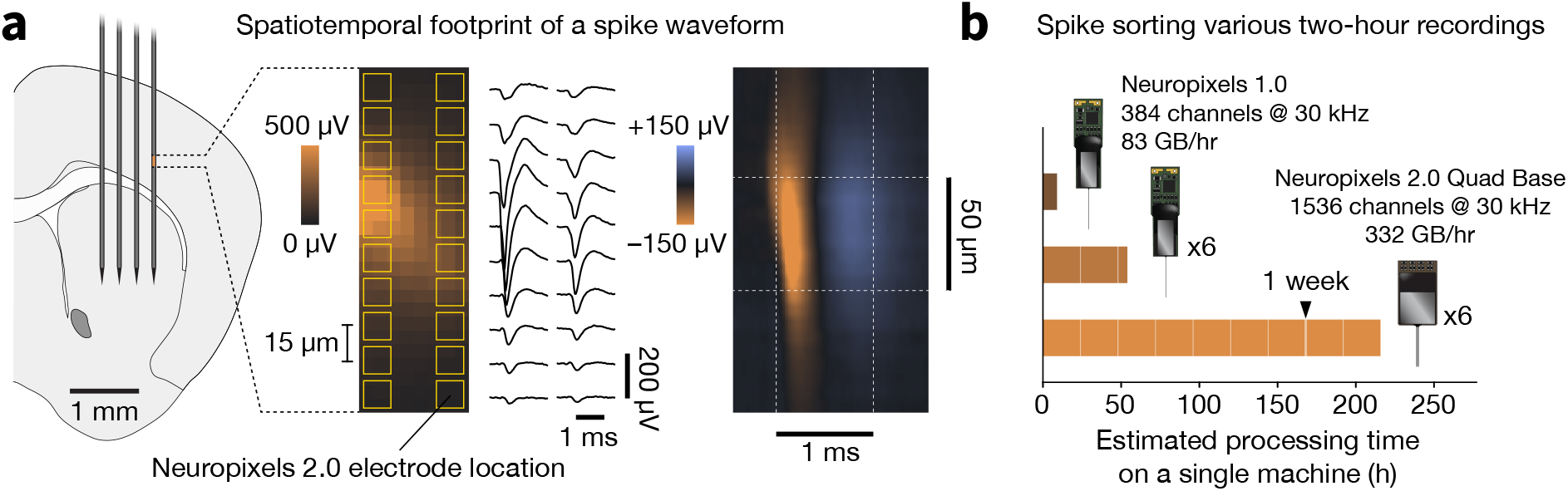
Challenges of scaling electrophysiological recordings. **a**, Multi-shank Neuropixels probe overlaid on a mouse brain, with zoomed-in region showing the spatial footprint of a typical spike waveform (from (Steinmetz & Ye 2022)). Waveforms from individual electrodes and spatiotemporal footprint (sampled by a hypothetical Neuropixels 2.0 probe) are shown on the right. The approximate scale of the spike (50 µm x 2 ms) necessitates dense sampling in both space and time. Scaling up the number of recorded neurons requires increasing the number of electrodes that are in close physical proximity to neurons. **b**, Times required to run preprocessing, spike sorting, and automated curation on two-hour recordings with different probe configurations. Assuming no parallelization across machines, a recording with six Neuropixels 1.0 probes (384 channels each) would take more than two days to process. A recording with six Neuropixels 2.0 Quad Base probes (1,536 channels each), which recently became commercially available, would take over one week. Parallelization is essential to complete processing in under 24 hours after data collection.

Because spike sorting is computationally intensive, benchmarking sorter performance is even more demanding, as it requires running multiple algorithms on the same underlying data. Many spike sorting algorithms have been optimized for large-scale electrophysiology data (Pachitariu et al. 2024; Yger et al. 2018; Chung et al. 2017; Boussard et al. 2023; Meyer et al. 2024), yet systematic comparisons of their suitability across brain regions, species, and electrode types remain scarce (Carlson & Carin 2019). Given the enormous investment in electrode technology, each step in the spike sorting process should be benchmarked to ensure we can extract the maximum value from our data.

To address the scaling challenge, we developed a core spike sorting pipeline designed for distribution across many workstations, either locally or in the cloud. This pipeline improves the efficiency of spike sorting individual experiments, reducing the estimated processing time for an experiment with six Neuropixels “Quad Base” probes (1,536 channels each) from over a week to just ten hours, a speedup of more than 20-fold. To enable more rigorous evaluation of spike sorter accuracy, we developed a benchmarking pipeline that runs variations of the core pipeline on real data with injected ground-truth spikes. Together, these pipelines are prepared to handle the coming onslaught of data from current and future electrophysiology device.

The implementation of our pipelines was facilitated by three established technologies: `Nextflow` (Di Tommaso et al. 2017), `SpikeInterface` (Buccino et al. 2020), and `Code Ocean` (Cheifet 2021). `Nextflow` provides an abstraction layer between the data processing steps and the specific resources needed to execute them, meaning there are minimal modifications required to run the pipeline on a single machine, a high-performance computing cluster, or the public cloud. `Nextflow` also enables modularity by allowing processing steps with unique hardware or software dependencies to be seamlessly integrated into the same workflow. `SpikeInterface` is a Python package that makes a wide range of validated algorithms for data preprocessing, spike sorting, and curation available via a unified API. `SpikeInterface` encapsulates each processing step into its own module, allowing users to mix and match different algorithms to suit their needs. `Code Ocean` is a cloud platform for reproducible scientific computing. It is not required for running our pielines, but it greatly simplifies the process of deploying them in the cloud and streamlines the transition to downstream analysis in a scientist-friendly development environment.

**Reproducibility** is critical for spike sorting, as small differences in software versions or parameters can lead to widely divergent results. Cloud deployment addresses this by running code in containerized environments, ensuring identical processing across datasets. Prior attempts to improve reproducibility by migrating spike sorting to the cloud have important shortcomings. SpyGlass (Lee et al. 2024) is a data management and analysis framework built on top of DataJoint (Yatsenko et al. 2018) that includes spike sorting capabilities. Although it is designed to be run in the cloud, it does not manage parallelization, which limits its **scalability** as channel counts increase. NeuroCAAS (Abe et al. 2022) allows neuroscientists to run predefined analyses in the cloud by dragging and dropping data files into a browser window. While this approach makes the barrier to entry extremely low, it lacks the **modularity** needed to readily swap in new slgorithms or chain together different combinations of processing steps. Geng and colleagues recently described a cloud-based pipeline optimized for high-density multielectrode arrays (Geng et al. 2024). Its reliance on custom Kubernetes infrastructure constrains **portability** and prevents straightforward deployment on alternative backends that are critical for most research groups.

Previous efforts to benchmark spike sorting algorithms have mainly relied on simulations of extracellular spikes, which are computationally intensive and often fail to capture the complexities of real data (Martinez et al. 2009; Buccino et al. 2020; Laquitaine et al. 2024; Hagen et al. 2015; Buccino & Einevoll 2021). Benchmarking with genuine ground truth data obtained from simultaneous intracellular and extracellular recordings provides the most realistic form of evaluation, but such datasets remain exceptionally scarce. SpikeForest (Magland et al. 2020) was a noteworthy attempt to aggregate and standardize benchmarking across available ground truth datasets, but it has not been updated to include more recent algorithms and therefore provides a limited and outdated view of the spike sorting landscape. An alternative strategy relies on hybrid benchmarking, in which groundtruth spikes are injected into real recordings. The paper describing `Kilosort4` used hybrid data to compare this algorithm to ten others (Pachitariu et al. 2024). However, the benchmarking framework they developed was not intended for extension to diverse recording conditions. SHYBRID (Wouters et al. 2021) offers a graphical interface for hybrid benchmarking and integrates with `SpikeInterface`, but does not incorporate cloud-based parallelization. Our approach integrates the innovations of our core spike sorting pipeline to enable practical benchmarking of hybrid large-scale electrophysiology datasets. Given the time-consuming nature of individual pipeline steps, parallelization allows us to systematically compare algorithms over the span of hours, rather than weeks.

In the sections that follow, we first describe the three underlying technologies that enabled us to build spike sorting pipelines that meet our requirements of reproducibility, scalability, modularity, and portability. We then provide an overview of our core spike sorting pipeline, which has already processed data from more than 1,000 multi-probe recordings. We then present our benchmarking pipeline, which we use to compare the performance of `Kilosort4` and `Kilosort2.5` and to assess the impact of lossy compression—a strategy that could greatly reduce data volumes but may compromise sorting accuracy. Taken together, these pipelines support our overarching aim: to more accurately and efficiently connect the activity of individual neurons with population-level dynamics that span the brain.

## Results

### Enabling technologies

Three existing software tools form the foundation for our pipelines:

`Nextflow` (Di Tommaso et al. 2017) is a domain-specific language for orchestrating scientific data processing pipelines. In `Nextflow`, a *process* is a self-contained unit that performs a specific task. Processes are linked together through *channels*, which handle data flow between tasks, ensuring clear input/output relationships.

`SpikeInterface` is an open-source Python package for processing extracellular electrophysiology data (Buccino et al. 2020). It includes several modules that encompass all aspects of extracellular electrophysiology data analysis, including reading data (from more than 30 file formats), preprocessing, spike sorting (with at least 10 different spike sorters), postprocessing, curation, visualization, and more.

`Code Ocean` (Cheifet 2021) is a cloud platform for reproducible scientific computing. `Code Ocean` was originally developed to support reliable regeneration of figures and analyses in journal publications using open source tools. The platform introduces the concept of the *capsule*, which integrates the three components necessary to fully reproduce a result: immutable data, a fully specified execution environment, and version-controlled code that can be run with a single command. `Code Ocean` has native support for `Nextflow` pipelines and makes them easy build and operate using scalable cloud resources. More generally, `Code Ocean` eases the transition to working with data in the cloud, providing a variety of familiar development environments (Visual Studio Code, JupyterLab, RStudio, MATLAB, and Ubuntu virtual desktops) with file-based access to data. Many scientific software tools only support traditional file-based data access patterns and would otherwise require extensive customization to handle cloud object storage APIs.

Leveraging these technologies was essential for meeting our design requirements of **reproducibility, scalability, modularity**, and **portability**:

- **Reproducibility** is a cornerstone of our pipelines. Each `Nextflow` process points to a specific image of a Docker or Singularity container, which guarantees that the software environment remains consistent across different computational backends. Each spike sorter supported by `SpikeInterface` ships with a container image available on `DockerHub`). This ensures the same version of the sorter can always be re-run, eliminates installation headaches, and simplifies deployment on cloud infrastructure. `Code Ocean` adds an additional layer of reproducibility by tracking all processing steps that happen upstream or downstream of our spike sorting pipeline. This feature is helpful when preparing figures for publication, when the details of the entire analysis chain (not just the spike sorting outputs) must be transparently shared.
- **Scalability** is achieved by leveraging distributed computing to support parallelization over multiple probes. Since each `Nextflow` process runs independently and communicates through channels, it enables seamless parallel execution of processes, making the workflow highly scalable. Furthermore, `Nextflow` can provision custom resources for each process, meaning that more expensive cloud instances with GPUs do not need to be deployed beyond the spike sorting step. This adaptability allows users to tailor their computational resources to the complexity and size of their datasets. In addition, `SpikeInterface` makes use of parallelization wherever possible, e.g. for filtering, compression, waveform extraction, and a range of other processing steps.
- **Modularity** is enforced by encapsulating each pipeline step into `Nextflow` processes and channels. This makes it simple to swap out algorithms or incorporate novel pre- and post-processing steps without changing the overall workflow. `SpikeInterface` also promotes modularity by defining standard formats for transferring data between processing steps.
- **Portability** is enabled by `Nextflow` *executors* that are compatible with a variety of backends. To simplify deployment on commonly used backends for academic institutions, such as multi-processor workstations and SLURM HPC systems, we provide pre-configured scripts, configuration files, and detailed documentation. For scientists interested in using our pipeline in the cloud, `Code Ocean` is by far the easiest way to get up and running, since it natively supports `Nextflow` over an Amazon Web Services (AWS) Batch backend. However, `Nextflow` workflows can also run on major cloud providers (including AWS, Google Cloud, and Microsoft Azure) with minimal configuration changes.

### An end-to-end pipeline for spike sorting large-scale electrophysiology data

We designed a pipeline for spike sorting electrophysiology data, ensuring reproducibility, scalability, modularity, and portability. The pipeline addresses critical aspects of data processing, from ingestion of raw data to the curation of spike sorting outputs. The spike sorting pipeline is publicly available on `GitHub` (AllenNeuralDynamics/aind-ephys-pipeline) and detailed documentation is hosted on `ReadTheDocs` (aind-ephys-pipeline.readthedocs.io).

The pipeline encompasses eight major steps (Figure 2):

**Figure 2:**
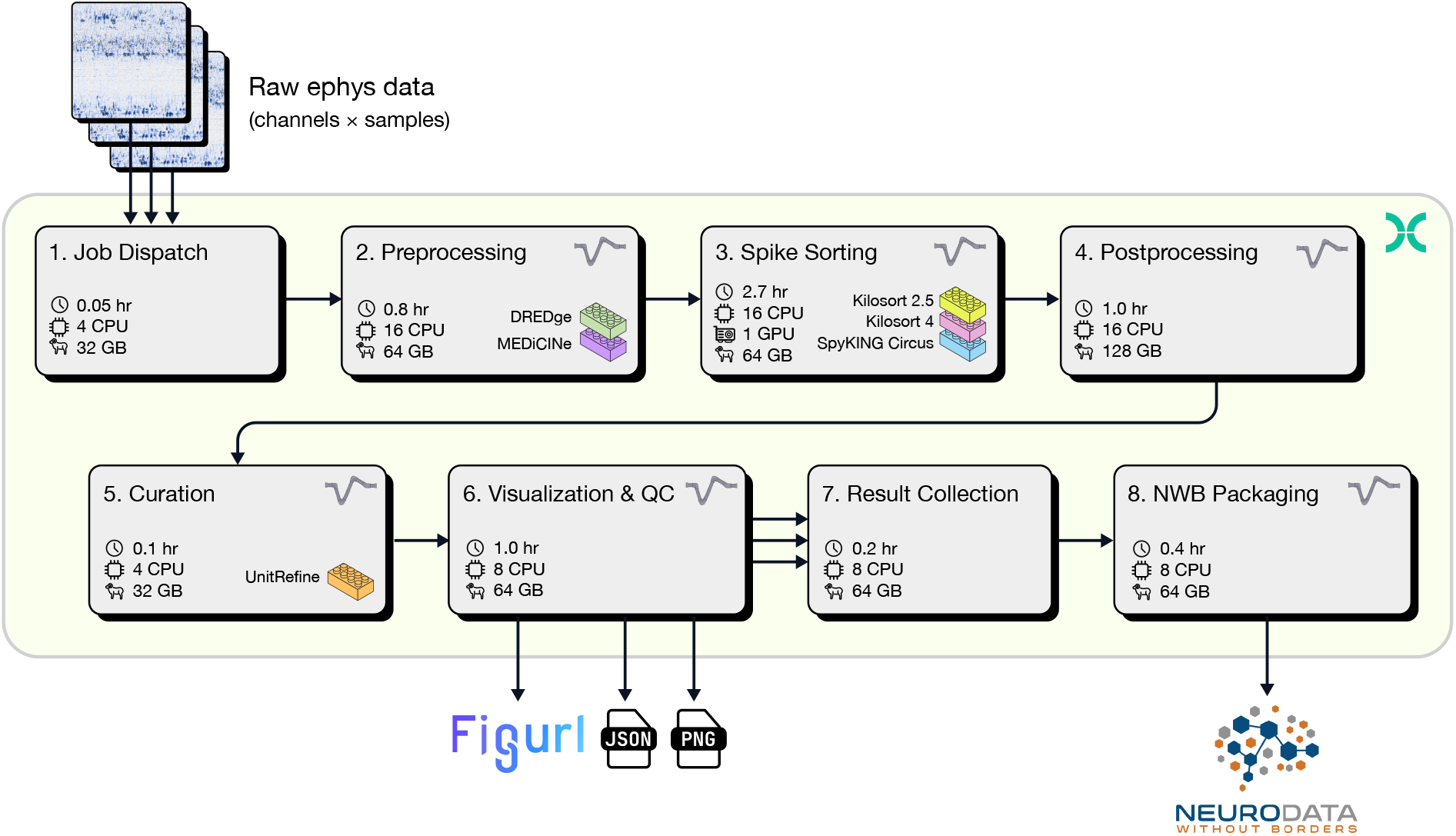
Spike sorting pipeline overview. Raw electrophysiology data from multiple probes (top) is ingested by the Job Dispatch step, which coordinates parallelization of downstream processing. All steps are run in parallel until Result Collection. Pipeline outputs (including `Figurl` interactive visualizations, metrics stored in JSON format, PNG-formatted images, and a Neurodata Without Borders file) are shown at the bottom. Each step includes an estimate of run time (per hour of recording) and required computing resources (CPU, GPU, and RAM). The pipeline is encapsulated in a `Nextflow` workflow (green background), and individual steps are implemented in `SpikeInterface` (action potential logo). Interlocking brick icons indicate processing algorithms that can be easily substituted. For the spike sorting step, run time is calculated for `Kilosort4`. See Table 1 for detailed run times and cost estimates.

#### 1. Job dispatch

The entry point of the pipeline is a *job dispatch* step, which handles data ingestion and orchestration of parallelization. This step parses the input folder containing data from one recording session, which may include an unlimited number of probes. It outputs a set of configuration files containing key metadata about each session and recording, as well as the information to instruct `SpikeInterface` how to reload the recording. A parallel instance of downstream processing is launched for each configuration file. Parallelization is performed across *streams* (e.g., individual probes), *groups* (e.g., shanks of the same probe), and *recordings* (e.g., segments of continuous data). As an example, for a session with data from three Neuropixels 2.0 multi-shank probes with three recordings each, the job dispatch will output 36 configuration files (3 probes × 4 shanks × 3 recordings), which will spawn 36 parallel downstream processes.

**Table 1:** Resources, run time, and cost for all steps in the spike sorting pipeline. Values are based on a one-hour recording with six Neuropixels probes. Values for non-parallel steps (Job Dispatch, NWB Packaging, Result Collection) are normalized by the number of probes. For steps that run in parallel, run times and estimated cost are reported as the mean *±* standard deviation for probes in the same session. *Run Time per hour of recording with one 384-channel probe. **Cost per hour of recording with one 384-channel probe. Costs are estimated using the prices of the reported instance from https://aws.amazon.com/ec2/pricing/on-demand/ as of April 30, 2025.

| Step | Parallel? | CPUs | GPUs | RAM (GB) | Run Time* (h) | Instance | Cost** (USD) |
| --- | --- | --- | --- | --- | --- | --- | --- |
| Job Dispatch | N | 4 | 0 | 32 | 0.05 | r5a.xlarge | 0.011 |
| Preprocessing | Y | 16 | 0 | 64 | 0.84±0.19 | m5a.4xlarge | 0.58 |
| Spike Sorting (KS4) | Y | 16 | 1 | 64 | 2.7±0.12 | g4dn.4xlarge | 3.24 |
| Postprocessing | Y | 16 | 0 | 128 | 1.0±0.17 | r5a.4xlarge | 0.9 |
| Curation | Y | 4 | 0 | 32 | 0.1±0.1 | r5a.xlarge | 0.022 |
| Visualization/QC | Y | 8 | 0 | 64 | 1.02±0.12 | r5a.2xlarge | 0.46 |
| NWB packaging | N | 8 | 0 | 64 | 0.38 | r5a.2xlarge | 0.17 |
| Result Collection | N | 8 | 0 | 64 | 0.16 | r5a.2xlarge | 0.072 |
| Total | - | - | - | - | 6.25 |  | 5.554 |

#### 2. Preprocessing

The *preprocessing* step prepares raw electrophysiology signals for spike sorting. Four computations are applied in sequence:

- **Phase-shift correction**: Some high-density recording devices, such as Neuropixels, have fewer analog-to-digital converters (ADCs) than recording channels. During each sampling period, each ADC digitizes voltages from multiple electrodes, a process known as “multiplexing.”. Sample times for different groups of channels are therefore offset in time by a known amount. The phase shift algorithm uses a fast Fourier transform to make the signals appear as though they were sampled simultaneously across all channels. This correction increases the effectiveness of the subsequent denoising step (International Brain Laboratory 2024). If the input recording does not require phase shift correction, this step is skipped.
- **Filtering**: By default, a highpass filter with cutoff frequency of 300 Hz is applied, to preserve high-frequency information in spike waveforms. Users can opt for a bandpass filter to better remove high-frequency noise.
- **Denoising**: This step first masks out noisy or dead channels, then applies a Common Median Reference (CMR) or a highpass spatial filter (also referred to as *destriping*). By default, CMR is used, since destriping can create artifacts in spike waveforms (International Brain Laboratory 2024).
- **Motion estimation/correction**: Drift of the electrodes relative to the brain tissue is common issue in recordings with long, needle-like probes. Among the several methods are available for estimating and correcting drift, `DREDge` is used by default (Windolf et al. 2025). The estimated motion signal is used for visual assessment of the severity of drift throughout the recording. It is also possible to use the estimated motion to correct the traces via interpolation. However, since the spike sorting step may also include its own motion correction algorithm, interpolation is not performed by default.

The preprocessing step does not currently include artifact detection and removal (similar to the `gfix` method implemented by `CatGT` from the `SpikeGLX` tool (Karsh, n.d.)), but a similar processing step will be included in a future release.

#### 3. Spike Sorting

*Spike sorting* identifies discrete spike times in continuously sampled voltage traces and assigns each one to a “unit,” which may include spikes from one or more neurons. Currently, three spike sorting algorithms are available within the pipeline: `Kilosort4`, `Kilosort2.5` (Pachitariu et al. 2024), and `SpyKING-CIRCUS2` (Yger et al. 2018) (a new version of this algorithm fully implemented within `SpikeInterface`). User can modify any sorter parameters via a configuration file.

#### 4. Postprocessing

*Postprocessing* combines the preprocessing and spike sorting outputs to perform additional computations relevant for visualization, curation, and downstream analysis. This step begins by estimating each unit’s *template* (average extracellular waveform) and *sparsity* (the set of channels the template is defined on). Next, duplicated units, that can arise when using template-matching methods if different templates are consistently fit to the same spikes, are removed based on the fraction of overlapping spikes.

Several additional postprocessing *extensions* are computed and saved to the results, including:

- *waveforms*: a set of subsampled spike waveforms (maximum 500 per unit)
- *spike amplitudes*: peak-to-peak amplitudes of each spike in µV
- *template similarity*: pairwise template similarity across units
- *correlograms*: auto- and cross-correlograms for all units
- *ISI histograms*: inter-spike-interval histograms
- *unit locations*: estimated unit locations via monopolar triangulation
- *spike locations*: estimated spike locations via grid convolution
- *template metrics*: a set of metrics describing the spatial and temporal profile of each template
- *quality metrics*: a set of metrics for assessing the completeness and contamination of each unit

A comprehensive explanation for these extensions can be found in the **Methods** section.

#### 5. Curation

The *curation* step applies labels to sorted units to automate, or at least speed up, unit inspection and curation. Two set of labels are applied. The first is based on a set of quality metric thresholds (Siegle et al. 2021). Units are tagged as passing a *default_qc* when they satisfy the criteria based on quality metric thresholds. Thresholds can be user defined, and these are the default:

- ISI violation ratio < 0.5 (places an upper bound on spike train contamination)
- Amplitude cutoff < 0.1 (removes units with a large fraction of spikes below the detection threshold)
- Presence ratio > 0.8 (removes units that were not detected for more than 20% of the recording duration)

The second set of labels is based on `UnitRefine` (Jain et al. 2025), which uses two pre-trained random forest classifiers to label units as noise, single-unit activity (SUA), or multi-unit activity (MUA).

#### 6. Visualization and quality control

The visualization step creates interactive views using `Figurl` (Magland & Soules 2025a) and `sortingview` (Magland & Soules 2025b). This technology, integrated into `SpikeInterface`, generates shareable links that can be viewed from any device. Our pipeline includes one link for visualizing the raw/preprocessed traces, drift maps, and estimated motion (Figure 3a) and one link for visualizing and curating the spike sorting results (Figure 3b). While we expect that machine learning tools will eventually automate the entire spike sorting process, we recognize that some manual curation may be beneficial in the present day. Therefore, the `sortingview` link can be used for merging and labeling units, the outputs of which can be incorporated into downstream analysis.

**Figure 3:**
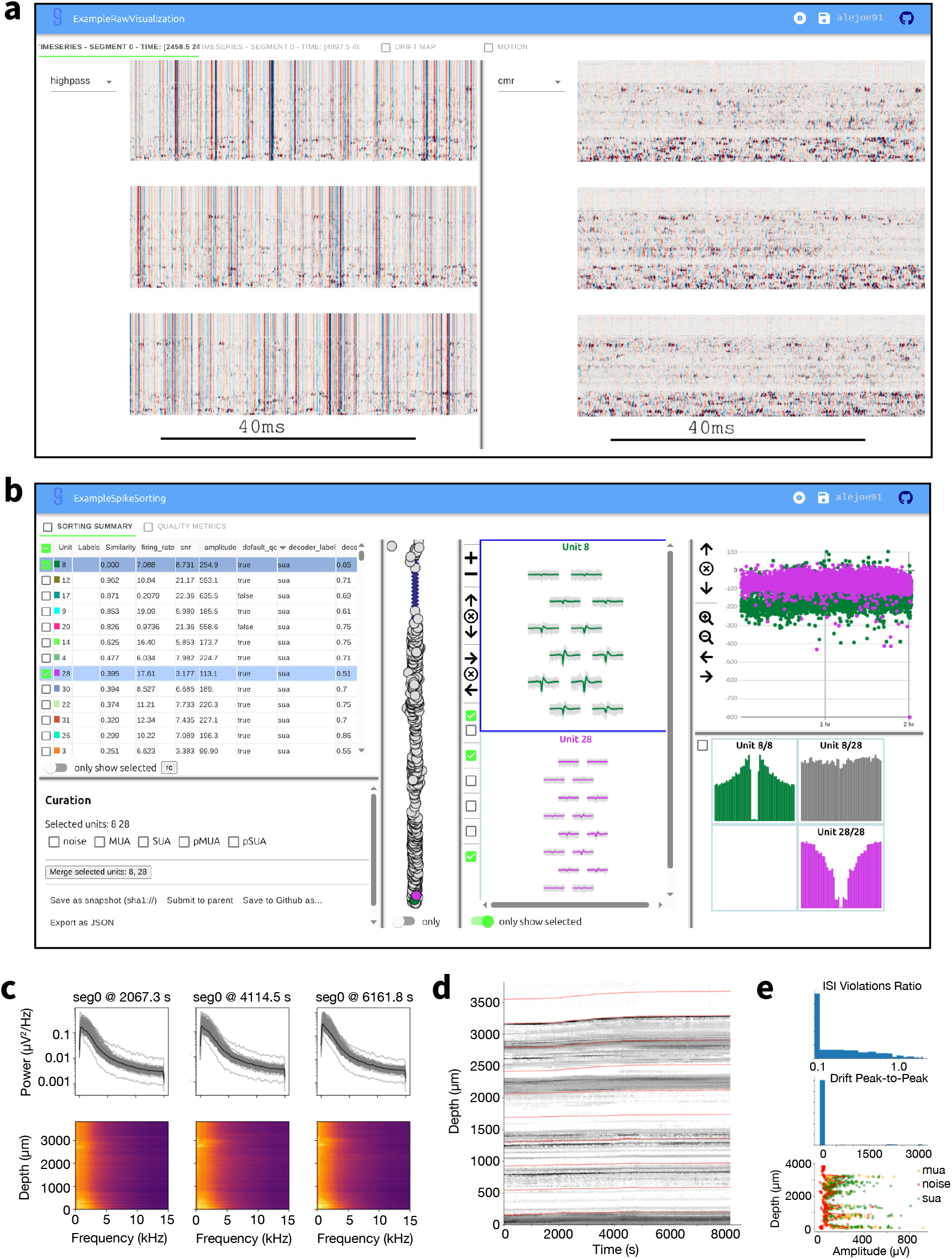
Outputs of the visualization and quality control step. **a**, Screenshot of interactive visualization of raw and preprocessed data (view on web). **b**, Screenshot of `SortingView` GUI, used for inspecting and curating spike sorting outputs (view on web). **c**, Quality control plot showing power spectra of continuously recorded electrical signals at three timepoints during the session. **d**, “Drift map” of spike locations over time, used to assess the overall stability of the recording. **e**, Histograms and scatter plot of key unit-specific quality metrics.

In addition to these shareable web-based visualization, the pipeline implements a quality control step that produces a series of static plots and metrics designed to asses the raw and processed data.

- Raw data: snippets of AP and LFP traces at multiple points over the session
- PSD: Power Spectrum Density of wide, high-frequency and low-frequency bands of each channel (Figure 3c)
- Noise: RMS channel values of raw data and preprocessed data
- Drift: raster map and estimated drift (Figure 3d)
- Saturation: count of positive/negative saturation events
- Unit Yield: distributions of unit metrics, amplitudes, and firing rates, along with a unit classification summary (Figure 3e)
- Firing Rate: population firing rate over the session

#### 7. Result Collection

Once all upstream processing steps are finished, the *result collection* step aggregates results from different probes into structured outputs, organizing them for downstream analysis and ensuring data integrity.

#### 8. NWB Packaging

The final step exports results into the widely used Neurodata Without Borders (NWB) format (Rübel et al. 2022). Each NWB file includes data from all probes that were recorded simultaneously. Prior to being shared, NWB files generated by this pipeline will need to be extended with subject metadata and behavioral information. We chose not to require these elements to generate the output files, so that the pipeline would only require raw electrophysiology data as input. However, if desired, subject and session information can be specified using additional input metadata files (following the Allen Institute for Neural Dynamics (AIND) Data Schema specification, see **AIND Data Schema**).

The pipeline supports multiple input data formats, including those from the most widely used applications for Neuropixels data acquisition (`SpikeGLX`, `Open Ephys`). Although the present work focuses on Neuropixels, our pipeline is not limited to them: the input layer supports any of the readers supported by `SpikeInterface` (over 30 file formats). We also provide the ability to read directly from an NWB file, for cases where raw data has been shared in this format.

The pipeline saves the outputs of all steps, which can be easily reloaded by `SpikeInterface` if additional processing is needed. In addition, the pipeline produces NWB files with spike times, waveforms, and metrics for all units. See **Spike sorting pipeline output** for more details.

Each step of the pipeline can be configured with several parameters. We provide a detailed list and description of parameters associated with each step as part of the pipeline documentation. Parameters can be passed to the pipeline either via a configuration file or via command line arguments. For example, the user can choose the type of filter (highpass/bandpass) or denoising strategy (CMR/destriping) or fine-tune individual spike sorting parameters, which are all exposed in the parameter file.

### Cost-effective and robust spike sorting

Table 1 reports the resources, run time, and estimated cost of each pipeline step. The effective run time for a specific input datasets can be calculated as the sum of non-parallel steps × duration × number of probes, plus the run time of the parallelized steps × duration. The cost is the total cost × duration × number of probes.

To provide a concrete example, for a two-hour session with six Neuropixels 1.0 probes and a total of 4563 spike sorted units, the effective run time is 18.40 hours^1^ and the total cost is 66.65 USD^2^. Although this may seem like a high cost for spike sorting a single experiment, it is negligible in comparison to the price of purchasing, installing, and maintaining a computing cluster with at least 6 GPU nodes. To appreciate the value of parallelization and targeted resource allocation, if the same spike sorting job ran sequentially on a single GPU-capable cloud workstation (`g4dn.4xlarge` 1.204 $/h), it would take approximately 75 hours (6.25 * 6 probes * 2 h) and cost over 90 USD. Therefore, job orchestration and parallelization not only reduce the overall run time by more than 4x, they also reduce the price by 27% (given the higher cost of GPU instances). Costs could be further reduced by choosing cheaper instances with fewer CPUs, at the expense of longer processing times.

Our spike sorting pipeline was first deployed on Code Ocean in February 2024, running `Kilosort2.5` as the default sorter. Since its initial deployment, it has processed an average of 17.2 sessions per week (Figure 4a). In November 2024, `Kilosort4` became the default spike sorter given its better performance (see **Benchmarking Application 1: Comparing** `Kilosort2.5` **and** `Kilosort4`). Several projects continued using `Kilosort2.5` for consistency. As of August 2025, the pipeline has processed a total of 1,287 sessions: 789 with `Kilosort2.5` and 468 with `Kilosort4` (Figure 4b). It is quite common in our sessions to record from more than one probe. The spike sorting pipeline has processed 3,126 individual probes (Figure 4c), yielding over 1 million individual units that were labeled as *neural* (or not noise) by `UnitRefine` and over 700’000 passing quality-metric-based thresholds (Figure 4d; see **Curation** section for more details).

**Figure 4:**
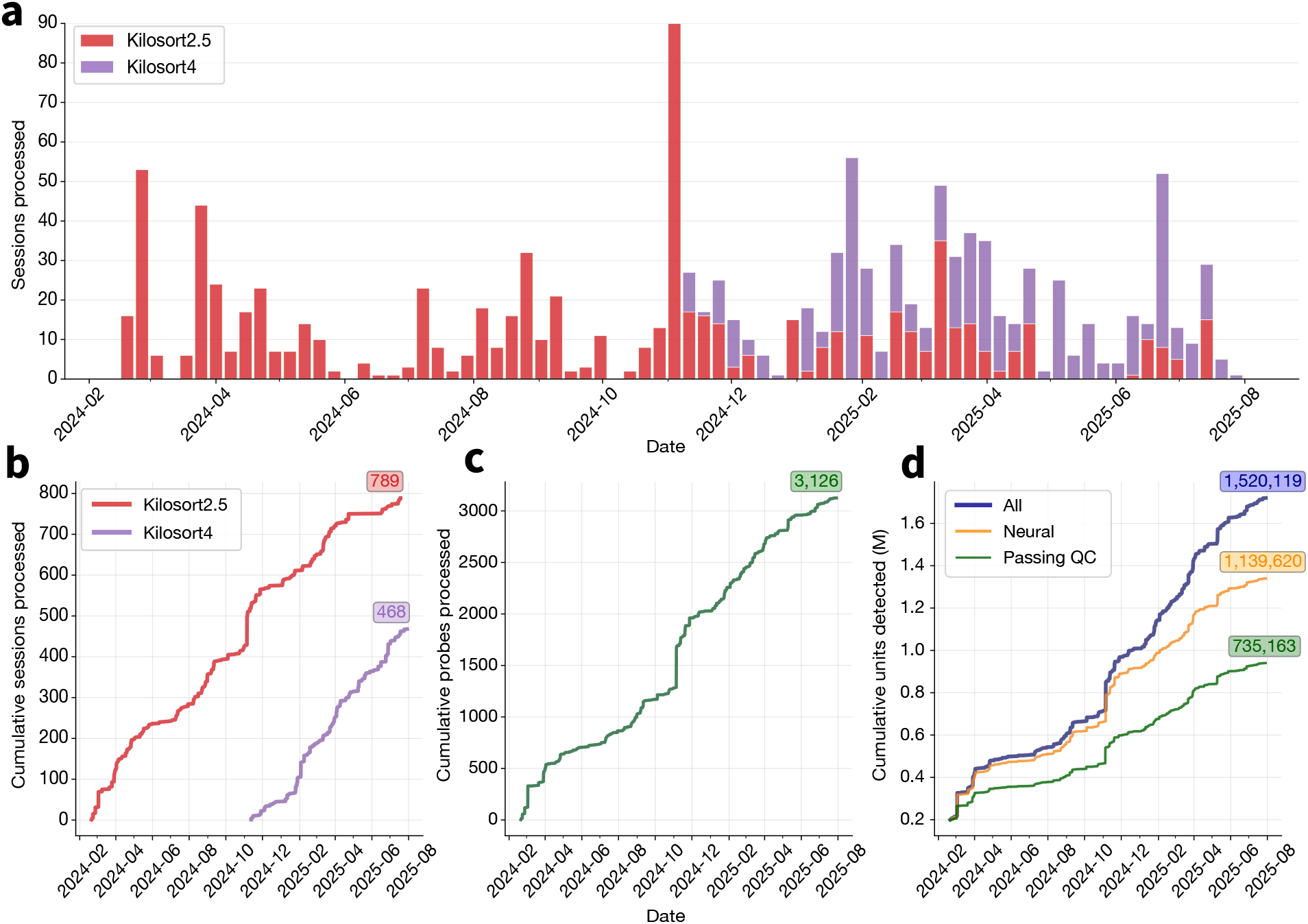
Spike sorting pipeline usage at the Allen Institute for Neural Dynamics. **a**, Sessions processed by the pipeline each week between February 2024 and August 2025. Color indicates the spike sorter used for each pipeline run. **b**, Cumulative sessions processed by the pipeline, split by spike sorter. **c**, Cumulative probes processed by the pipeline (many sessions include data from multiple probes). **d**, Cumulative units detected by the pipeline, with the subset of neural units indicated in yellow and units passing default quality metric thresholds in green. Units are considered neural if the `UnitRefine` classifier does not label them as “noise” (see **Curation** section for more details).

### A pipeline for rapid benchmarking of spike sorting algorithms and data processing steps

The same principles and technologies behind our spike sorting pipeline were used to develop a versatile pipeline for spike sorting evaluation. Evaluating spike sorting results requires access to ground truth spike times, so one can estimate what fraction of spikes are missed or improperly labeled. Real ground truth data, in which one has access to the actual spike times of a neuron recorded with an extracellular electrode, is limited to a few hard-won datasets, encompassing no more than a few dozen neurons (Henze et al. 2000; Neto et al. 2016; Yger et al. 2018; Allen et al. 2018; Marques-Smith et al. 2018). Therefore, we chose to base our evaluation pipeline on hybrid data, in which known spike templates are superimposed on real recordings. Hybrid data is a powerful tool for evaluating spike sorting options, since it maintains the inherent complexities of the data (e.g., spike correlations, noise statistics, and drift) (Rossant et al. 2016; Pachitariu et al. 2016; Pachitariu et al. 2024), while still providing the ground truth spike times needed to assess sorter accuracy. However, to avoid unwanted changes to the properties of the underlying data, a limited number of artificial spike trains should be superimposed on each original recording. This means that many iterations of spike sorting may be required to uncover statistically significant differences between processing steps.

We used the recently developed hybrid data generation framework in `SpikeInterface` to construct a `Nextflow` pipeline for evaluating spike sorting options. As with the core spike sorting pipeline, `Nextflow` enabled straightforward parallelization of pipeline steps, reducing the effective run times by over two orders of magnitude in comparison to serial execution (Table 2). The full pipeline is available on `GitHub` (github.com/AllenNeuralDynamics/aind-ephys-hybrid-benchmark).

**Table 2:** Benchmarking pipeline run times. The number of jobs and effective run time in hours for the two benchmarking applications (`Kilosort2.5` vs `Kilosort4`, Lossless vs Lossy) on data from two types of Neuropixels probes (NP1, NP2). The “distributed” execution refers to our `Nextflow` implementation, while the “serial” execution has all jobs running sequentially on an individual workstation. The run time for serial execution is extrapolated from the distributed run times, and was not actually measured.

| Benchmark application | Probe type | Execution | # Jobs | Effective run time (h) |
| --- | --- | --- | --- | --- |
| Kilosort2.5 vs Kilosort4 | NP1 | distributed | 398 | 14 |
| Kilosort2.5 vs Kilosort4 | NP1 | serial | 398 | 870 |
| Kilosort2.5 vs Kilosort4 | NP2 | distributed | 662 | 12 |
| Kilosort2.5 vs Kilosort4 | NP2 | serial | 662 | 846 |
| Lossless vs Lossy | NP1 | distributed | 578 | 18 |
| Lossless vs Lossy | NP1 | serial | 578 | 1590 |
| Lossless vs Lossy | NP2 | distributed | 962 | 16 |
| Lossless vs Lossy | NP2 | serial | 962 | 1684 |

As input to the pipeline, we used recordings from commercially available single-shank Neuropixels 1.0 (Jun et al. 2017) and four-shank Neuropixels 2.0 (Steinmetz et al. 2021) probes. These probes not only differ in their site geometry, but also in their signal bandwidth and bit depth. Neuropixels 1.0 probes have a built-in 300 Hz high-pass filter, and use 10-bit analog-to-digital Converters (ADCs), while Neuropixels 2.0 probes acquire wideband signals with 12-bit ADCs. We processed each shank separately, for a total of 6 sessions × 6 probes = 36 Neuropixels 1.0 recordings and 15 insertions × 4 shanks = 60 Neuropixels 2.0 recordings. For each recording, we randomly injected 10 hybrid units across 5 instances, generating a grand total of 1,800 ground truth units for Neuropixels 1.0 and 3,000 for Neuropixels 2.0.

For all results, we perform spike train comparisons and compute performance metrics as defined in (Buccino et al. 2020), using all units returned by the spike sorter (without any sorter-specific curation). In brief, we match sorted spike trains to ground-truth ones based on their agreement (fraction of matched spikes within 0.2 ms). For each matched pair, we count the number of true positives (#TP: ground-truth spikes found by the sorter), false negatives (#FN: ground-truth spikes not found by the sorter), and false positives (#FP: sorted spikes not found in the ground truth). We then evaluated each matched pair based on the following metrics:

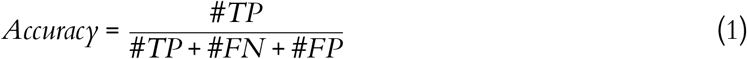

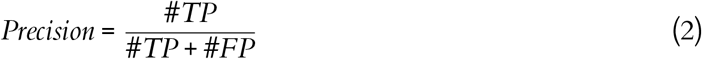

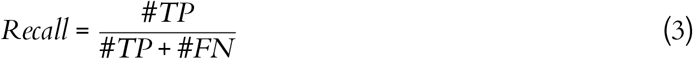

#### Pipeline steps

The benchmarking pipeline includes four major steps (Figure 5):

**Figure 5:**
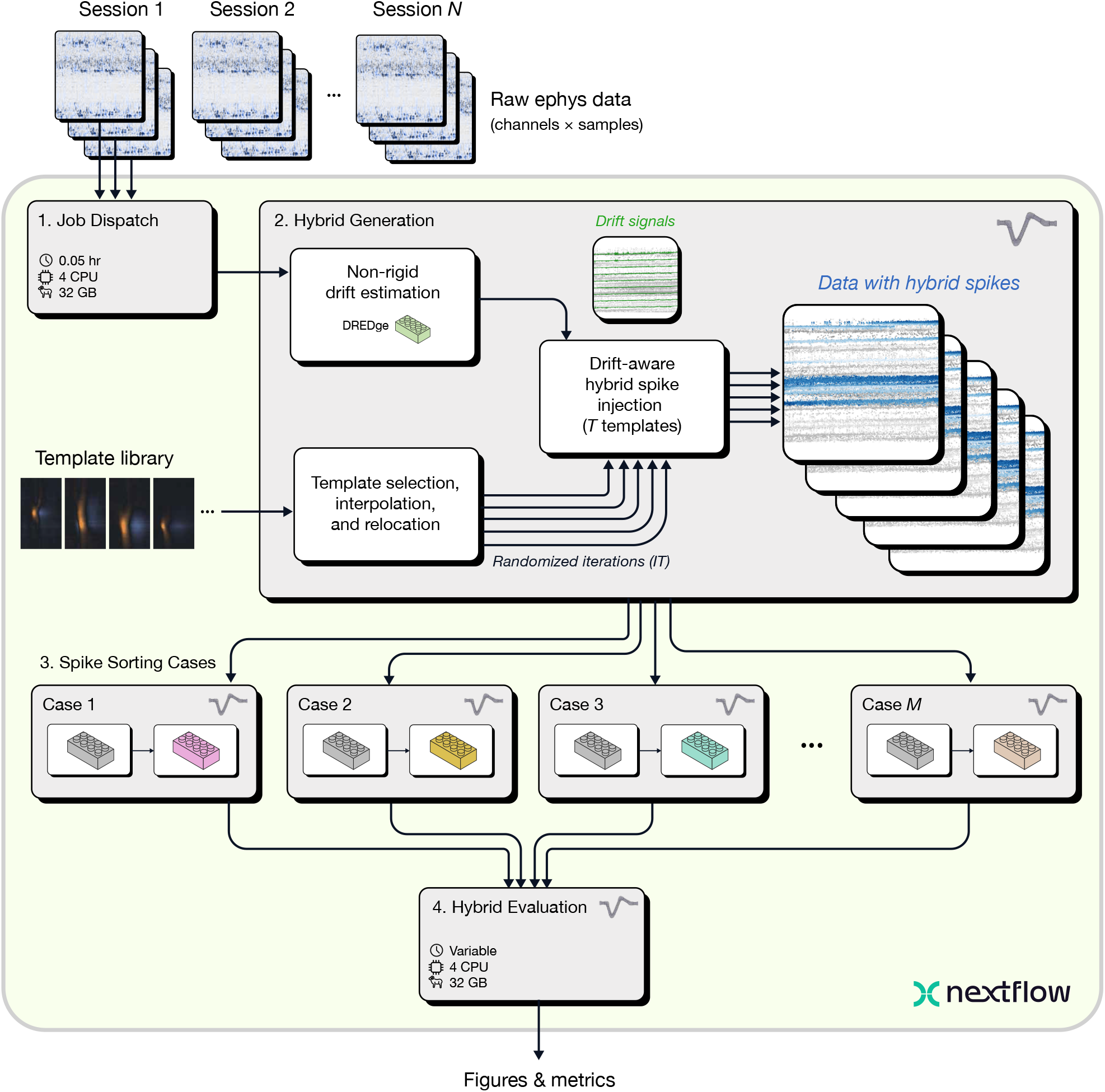
Benchmarking pipeline overview. The Job Dispatch step accepts *N* sessions as input, each of which may contain data from multiple probes. Next, the Hybrid Generation step injects *T* ground truth spike templates for each session and probe (default = 10), with *IT* randomized iterations (default = 5). Hybrid data are then processed in parallel through *M* Spike Sorting Cases, each of which sorts data using a different set of algorithms or parameters. The sorting results are then collected and compared by the Hybrid Evaluation step, which outputs figures and metrics for analyzing the performance of each case.

##### 1. Job Dispatch

The first step of the benchmarking pipeline is the same as that of the spike sorting pipeline: data ingestion and parallelization. The same input data formats are supported, but the benchmarking pipeline is capable of processing multiple sessions at a time to facilitate rapid evaluation across many input datasets in parallel.

##### 2. Hybrid Generation

For each input dataset, multiple hybrid recordings are generated by randomly sampling templates from a library available through `SpikeInterface`. The template library includes both Neuropixels 1.0 templates from the International Brain Laboratory (IBL) Brain-Wide Map (International Brain Laboratory et al. 2024; International Brain Laboratory et al. 2025) and high-density Neuropixels Ultra templates from (Steinmetz & Ye 2022). Here, we selected IBL templates for Neuropixels 1.0 probes and Neuropixels Ultra ones for Neuropixels 2.0, which were interpolated to the Neuropixels 2.0 geometry and relocated to cover the entire length of each shank. By default, 10 hybrid units, following a Poisson process with mean firing rates of 15 Hz, are added to each probe or shank, after rescaling to match a user-defined range (e.g., 50-200 µV). It is important for injected spikes to follow a recording’s natural drift profiles so that they smoothly blend in with the original spikes. The hybrid data generation step estimates non-rigid motion using `DREDge` (Windolf et al. 2025) and uses the motion signals to interpolate the selected templates in space during spike injection.

##### 3. Spike Sorting Cases

The generated hybrid recordings are processed in parallel through two or more *spike sorting cases* which can be specific to different applications. The spike sorting case is a subworkflow that can include multiple processes. Spike sorting cases can be used to compare different spike sorting tools, parameter sets of the same spike sorter, pre-processing options, or data compression algorithms.

##### 4. Hybrid Evaluation

The final step of the pipeline collects the spike sorting results of all spike sorting cases for each input session, evaluates their performance against hybrid ground-truth, and generates reports and tables for additional downstream analysis. In addition, results from all sessions are also aggregated into global results and plots.

The benchmarking pipeline accepts the same input types as the spike sorting pipeline, but it can take multiple sessions as input. This makes it straightforward to run the pipeline across many recordings without the need to modify the pipeline script or configuration. When using multiple input sessions, the evaluation step will aggregate outputs from all sessions into coherent results and visualizations.

The benchmarking pipeline outputs figures (as PNG images) and tables (as CSV files). In addition, all computed spike sorting outputs and comparisons are stored and can be reloaded by `SpikeInterface` if needed. More details can be found in the **Evaluation pipeline output** section.

The benchmarking pipeline reuses several capsules from the spike sorting pipeline. The supported input data formats and preprocessing parameters are therefore the same and can be specified similarly to the spike sorting pipeline presented above. The step that generates hybrid recordings takes the number of hybrid units per recording (default = 10) and the number of random iterations for each recording (default = 5) as additional inputs.

### Benchmarking Application 1: Comparing Kilosort2.5 and Kilosort4

Our first application of the benchmarking pipeline was comparing the performance of different spike sorting algorithms. We focused on `Kilosort`, the most widely used spike sorter for Neuropixels probes. The `Kilosort` algorithm has evolved alongside Neuropixels technology, with the initial version developed to efficiently sort data from Neuropixels 1.0 probes (Pachitariu et al. 2016). `Kilosort2.5` was introduced in the publication describing Neuropixels 2.0 probes (Steinmetz et al. 2021), and was incorporated into the original version of our spike sorting pipeline. `Kilosort3` included a new clustering algorithm, but for various reasons did not become as widely adopted as `Kilosort2.5`. `Kilosort4` was launched in February 2024 alongside benchmarking results showing that it outperformed all previous versions of `Kilosort` on hybrid datasets generated with Neuropixels 1.0 templates (Pachitariu et al. 2024). `Kilosort4` has many methodological and implementation differences with respect to `Kilosort2.5` (including being written in Python, rather than Matlab), so here we consider them two different spike sorters, rather than two versions of the same sorter.

For this application, the pipeline included two spike sorting cases:

1. preprocessing → `Kilosort2.5`
2. preprocessing → `Kilosort4`

Figure 6 provides an interpretable visual representation of the results for one Neuropixels 1.0 dataset. Figure 6a displays a zoomed in 100 ms snippet of traces with highlighted hybrid spikes. The red and purple spikes are found only by `Kilosort2.5` and `Kilosort4`, respectively. The blue spikes are found by both sorters. When we extend the view to the entire recording in the form of a spike raster (Figure 6b), it is clear that `Kilosort4` finds a higher proportion of hybrid spikes, with only a small number of spikes detected exclusively by `Kilosort2.5`. There was no clear degradation in terms of false positive (Figure 6c) and false negative rates (Figure 6d) for `Kilosort4`.

**Figure 6:**
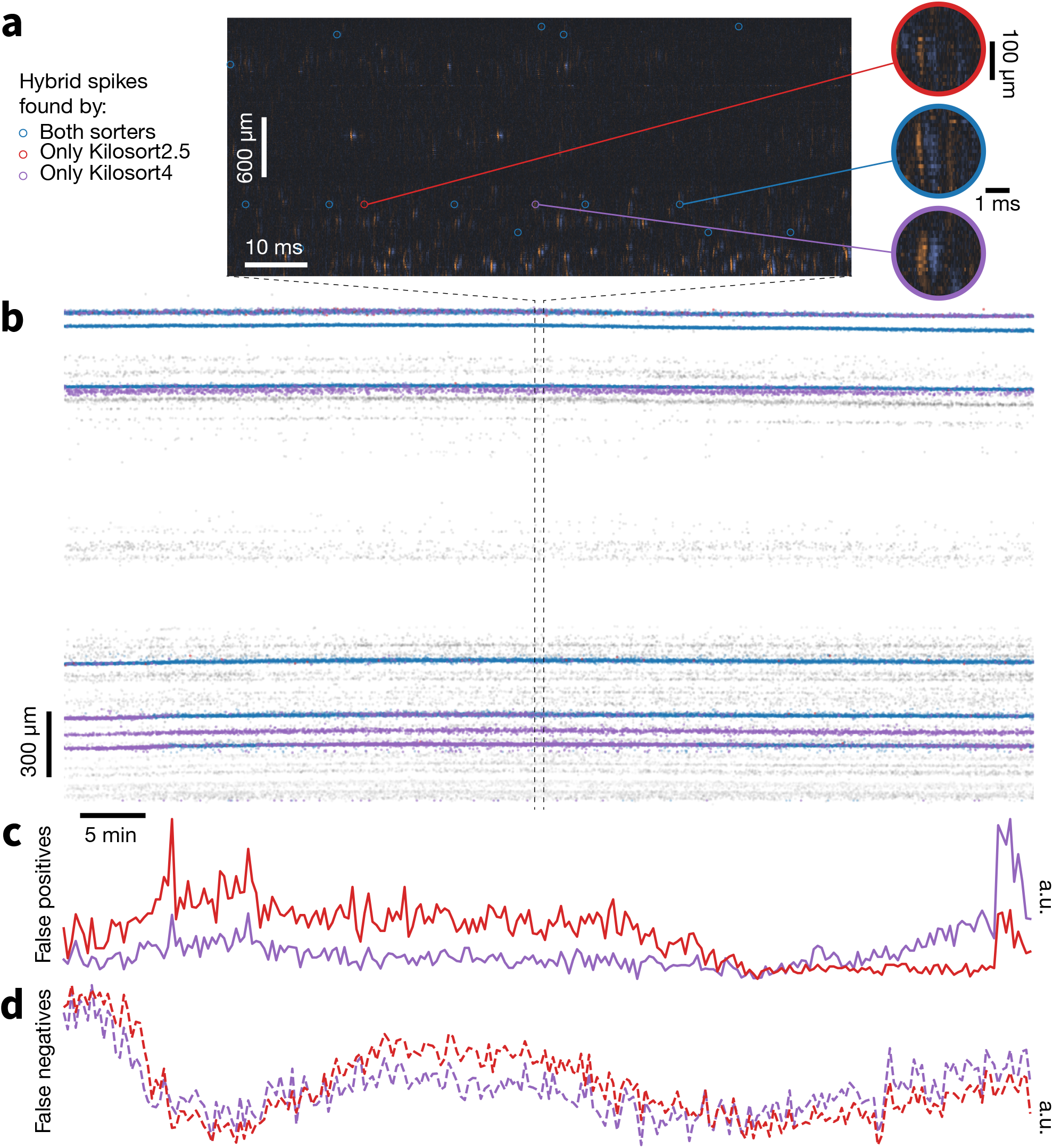
Example hybrid evaluation. **a**, 100 ms snippet of traces, showing ground-truth hybrid spikes and respective waveforms in *blue* if found by both sorters, *red* if found by `Kilosort2.5` only, and *purple* if found by `Kilosort4` only. **b**, Spike rasters of all hybrid spikes in the recording with same color scheme as panel a. Gray spikes are those from the original recording. Note that spikes not detected by any sorter are not visible in the plot. **c**, Fraction of false positive spikes found by either sorter (a.u. = arbitrary units). **d**, Fraction of false negative spikes. False positive and false negative spikes are calculated using 10 s time bins.

Across all recordings with both probe types, `Kilosort4` is able to find hybrid units with significantly greater accuracy (NP1: *P* < 1×10^−10^, effect size = 0.276; NP2: *P* < 1×10^−10^, effect size = 0.408), precision (NP1: *P* < 1 × 10^−7^, effect size = 0.272; NP2: *P* < 1 × 10^−10^, effect size = 0.232), and recall (NP1: *P* < 1 × 10^−10^, effect size = 0.319; NP2: *P* < 1 × 10^−10^, effect size = 0.415) (Figure 7a). These differences are most pronounced for units with signal-to-noise ratios below 10, which are generally more difficult to detect (Figure S1). When comparing individual ground truth units, `Kilosort4` tends to have higher accuracy and recall, at the expense of a slight degradation of precision (Figure 7b). This suggests that `Kilosort4` recovers more complete spike trains, but also finds more false positive spikes in the process. This can also be seen when looking at the distributions of the refractory period contamination (Llobet et al. 2022) for ground truth units detected by each sorter: `Kilosort4` has slightly higher values than `Kilosort2.5` (*P* < 1 × 10^−10^, effect size = 0.013)(Figure 7c). When looking at presence ratio, `Kilosort4` finds significantly more units which are active throughout the entire recording (*P* < 1 × 10^−10^, effect size = 0.193) (Figure 7d). This could be due to a more effective merging strategy implemented in `Kilosort4` (Pachitariu et al. 2024). Overall, these results demonstrate a clear superiority of `Kilosort4` in terms of spike sorting performance, although it does take `Kilosort4` almost twice as long to run as `Kilosort2.5` (Figure S2).

**Figure 7:**
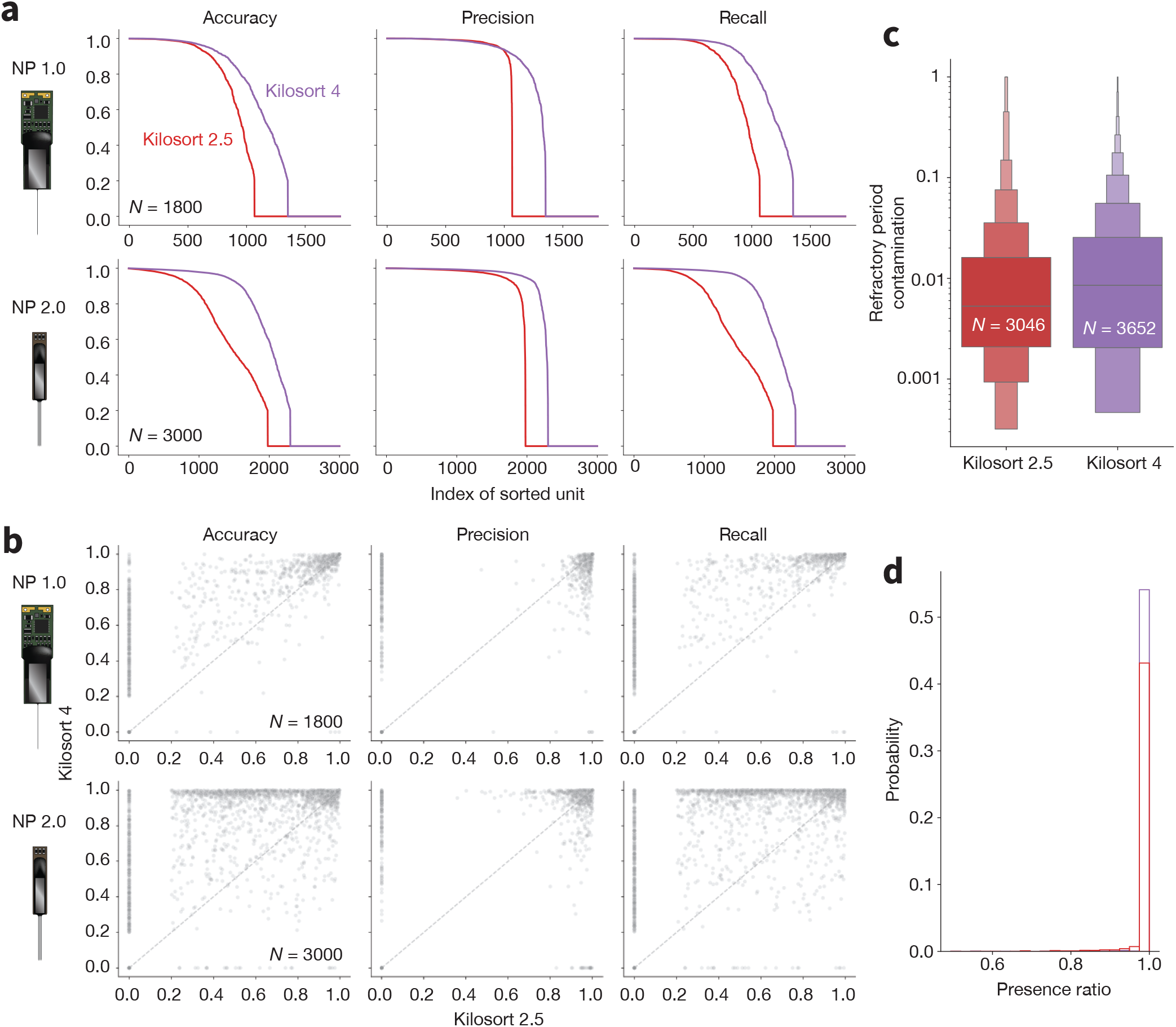
Benchmarking spike sorting algorithms. **a**, Accuracy, precision, and recall for all ground truth units for `Kilosort2.5` (red) and `Kilosort4` (purple) for Neuropixels 1.0 (top panels) and 2.0 (bottom panels) probes. **b**, Scatter plots for accuracy, precision, and recall for ground-truth units matched by both `Kilosort2.5` (x axis) and `Kilosort4` (y axis) for Neuropixels 1.0 (top panels) and 2.0 (bottom panels) probes. **c**, Distributions of refractory period contamination for all units found by both sorters. **d**, Histograms of presence ratio values for all units found by both sorters. In panels c and d, data from both probes are combined and include all units that were matched to a ground-truth hybrid unit (accuracy*≥*0.2 - `Kilosort2.5`: 3,046 matches; `Kilosort4`: 3,652 matches).

Table 2 displays the total number of jobs for this benchmark application and compares the estimated wall-clock run times for a single workstation and our distributed `Nextflow` implementation. The Neuropixels 1.0 benchmark ran 398 jobs, with a distributed effective run time of around 14 hours. For the Neuropixels 2.0 benchmark, the number of jobs increased to 662 (as individual shanks are processed separately) and the run time was of 12 hours. Running the same benchmarks on single workstations would have taken 870 hours (∼ 36 days) for Neuropixels 1.0 and 846 hours (∼ 35 days) for Neuropixels 2.0, highlighting the need for distributed pipelines for efficient evaluations.

### Benchmarking Application 2: Comparing lossless and lossy compression

We used our benchmarking pipeline to investigate whether lossy compression degrades spike sorter performance. In a previous study, we evaluated a wide range of compression algorithms and found that `WavPack`, a widely used audio compressor, is the best option for lossless compression of extracellular electrophysiology data (Buccino et al. 2023). `WavPack` also comes with a lossy mode, which compresses data down to a target average number of bits per sample (BPS). While we previously found no spike sorting degradation after lossy compression using simulated datasets (Buccino & Einevoll 2021), here we extended the analysis to hybrid datasets.

We compared lossless compression with `WavPack` against three different bits per sample values (from less to more lossy): BPS=3 (∼18.75% original file size, 5.33× compression ratio), BPS=2.5 (∼15.6% original file size, 6.4× compression ratio), and BPS=2.25 (∼14.06% original file size, 7.1× compression ratio, and the lowest supported BPS value). For comparison, the lossless compression ratio for Neuropixels 1.0 data was 3.53× (28.39 ± 1.02% original file size, *N* = 36 insertions); the lossless compression ratio for Neuropixels 2.0 data was 3.51× (28.51 ± 0.93% original files size, *N* = 15 insertions). Lossy compression thus cuts file sizes roughly in half, which has the potential to greatly reduce data storage costs (see (Buccino et al. 2023) for an in-depth cost analysis).

For this application, the evaluation pipeline included four spike sorting cases:

1. original data (lossless) → preprocessing → `Kilosort4`
2. `WavPack` compression (BPS=3) → preprocessing → `Kilosort4`
3. `WavPack` compression (BPS=2.5) → preprocessing → `Kilosort4`
4. `WavPack` compression (BPS=2.25) → preprocessing → `Kilosort4`

In all cases, compression was applied before any preprocessing took place. For Neuropixels 1.0, we compressed the AP stream only. For Neuropixels 2.0, we compressed the full wide-band data.

For Neuropixels 1.0 data, we observed no significant changes in the performance curves across the four compression cases (Figure 8a, top). Median accuracy, precision, and recall across all hybrid units stay roughly constant as compression ratio increases (Figure 8b, top). When looking at the unit-wise difference in performance between the lossless result and the BPS=3 case (Figure 8c, top), it is clear that units from the lossless sorting case tend to have slightly higher accuracy and recall, but these differences are typically small.

**Figure 8:**
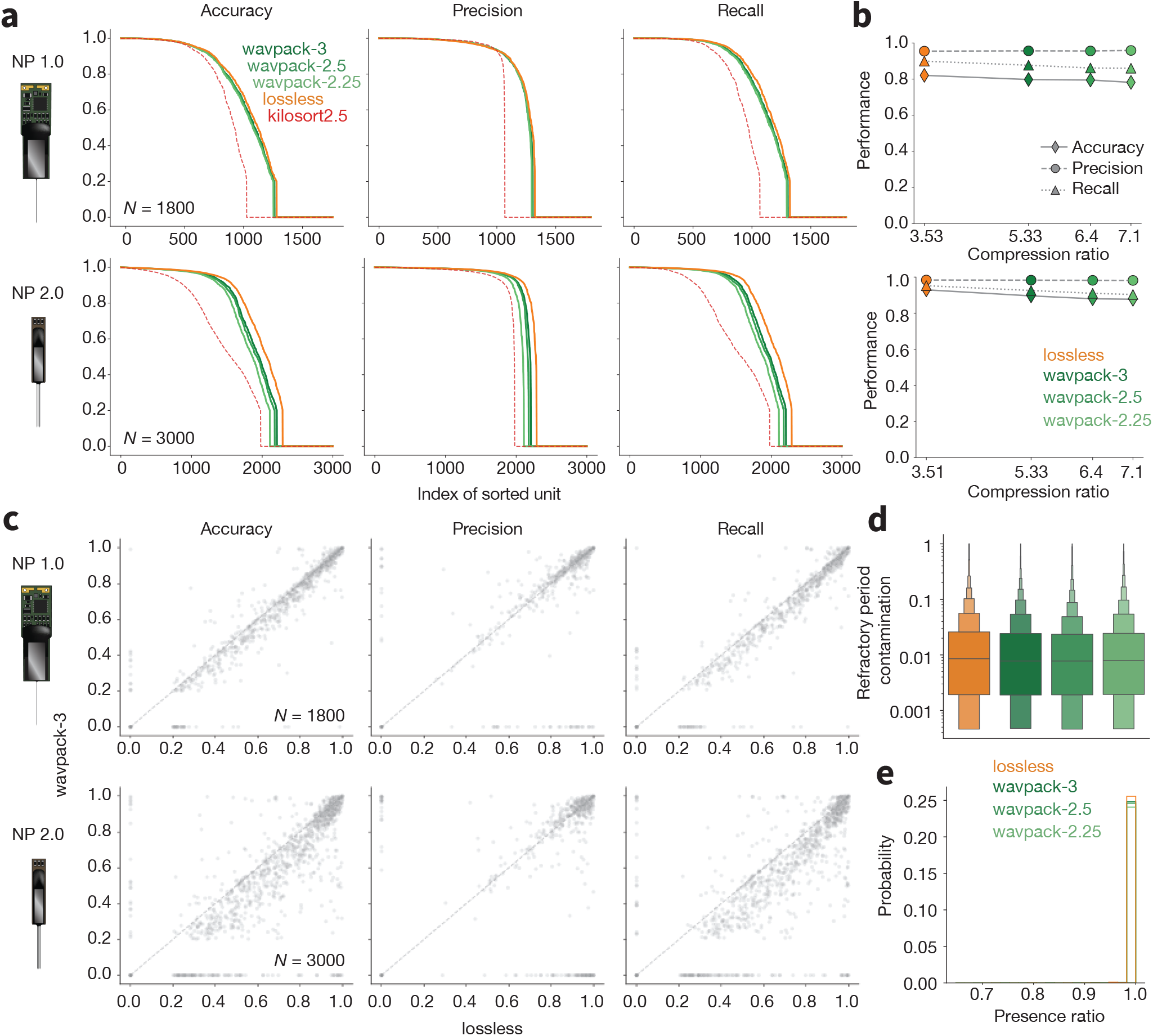
Lossy compression benchmarks. **a**, Accuracy, precision, and recall for all ground truth units for lossless, and lossy (dark green) `WavPack` compression (lighter greens as BPS decreases, i.e., compression ratio increases) for Neuropixels 1.0 (top panels) and 2.0 (bottom panels) probes. The dashed red line shows the curves for `Kilosort2.5` for reference. **b**, Median accuracy, precision, and recall as a function of compression ratio, for both probe types (top row: Neuropixels 1.0; bottom row: Neuropixels 2.0). **c**, Scatter plots for accuracy, precision, and recall for ground-truth units matched by both the lossless (x axis) and BPS=3 (y axis), for both probe types. **d**, Distributions of refractory period contamination for all units found at each compression ratio. **e**, Histograms of presence ratio values for all units found at each compression ratio. In panels d and e, data from both probes are combined and include all units that were matched with accuracy ≥ 0.2 to a ground-truth hybrid unit (Lossless: 3,612 matches; BPS=3: 3,513 matches; BPS=2.5: 3,475 matches; BPS=2.25: 3,405 matches).

The results for Neuropixels 2.0 show slightly more degradation of spike sorting accuracy with respect to the lossless compression results (lossless vs. BPS=3: *P* < 1 × 10^−10^, effect size = 0.08; lossless vs. BPS=2.5: *P* < 1 × 10^−10^, effect size = 0.1; lossless vs. BPS=2.25: *P* < 1 × 10^−10^, effect size = 0.13), precision (lossless vs. BPS=3: *P* < 1 × 10^−10^, effect size = 0.06; lossless vs. BPS=2.5: *P* < 1 × 10^−5^, effect size = 0.08; lossless vs. BPS=2.25: *P* < 1 × 10^−10^, effect size = 0.11), and recall (lossless vs. BPS=3: *P* < 1 × 10^−10^, effect size = 0.08; lossless vs. BPS=2.5: *P* < 1 × 10^−10^, effect size = 0.1; lossless vs. BPS=2.25: *P* < 1 × 10^−10^, effect size = 0.13) (Figure 8a - bottom) and can be mainly attributed to lower performance for low-SNR units (Figure S3b). For reference, Figure 8a also displays the `Kilosort2.5` performance curves in dashed red lines. This demonstrates that running `Kilosort4` on data with the highest level of lossy compression (BPS=2.25) still outperforms its predecessor on lossless data^3^.

For all well-detected units across both probe types, there were minimal differences in the refractory period contamination (Figure 8d, *P* = 0.044) and presence ratio (Figure 8e, *P* = 0.65). This indicates that, even for the highest compression settings, the overall quality of the detected units remains the same.

This benchmarking application had an effective run time of 18 hours for Neuropixels 1.0 and 16 hours for Neuropixels 2.0 (Table 2). Without distributed computing, the same evaluations would have taken 1590 hours (over 66 days) for Neuropixels 1.0 and 1684 hours (over 70 days) for Neuropixels 2.0.

## Discussion

We developed two pipelines for spike sorting large-scale electrophysiology data. The first encompasses all steps needed to extract spiking information from raw voltage traces and package them in a format that can be ingested for downstream analysis. The second is designed for rapidly evaluating the performance of different spike sorting algorithms, or measuring the impact of particular preprocessing steps on spike sorting accuracy. Both pipelines can run in the cloud, making it straightforward to parallelize almost every data processing step across multiple machines. When used in conjunction with `Code Ocean`, cloud resources can be provisioned via a user-friendly graphical interface, greatly lowering the barrier for entry for scientists who may have no prior experience with cloud computing. Importantly, though, the use of `Nextflow` for workflow orchestration means the same pipeline can also be deployed on local workstations or high-performance computing clusters in cases where these are preferred.

The pipelines balance the often competing demands of flexibility and reproducibility. The pipeline components are from `SpikeInterface`, because this package defines standardized Python objects for passing data between modules, making it straightforward to add, remove, or modify processing steps on demand. `SpikeInterface` (in conjunction with the `Neo` library) ensures the pipelines support input data from with virtually any acquisition device and recording apparatus used in the field. As new spike sorting tools and/or processing techniques are added to the `SpikeInterface` package, they will require minimal effort to be incorporated into our pipeline. As an example, the pipeline already supports two motion correction algorithms, DREDge (Windolf et al. 2025) and MEDiCINe (Watters et al. 2025), which were released in 2025. Maintaining this level of flexibility does not compromise reproducibility, as all code used in the pipeline can be traced to a specific commit in a `GitHub` repository, and all functions are executed inside standardized Docker or Singularity containers. This ensures that the exact same steps can be re-run in a reproducible manner, even if the hardware backend changes.

The spike sorting pipeline has been in use by all electrophysiology projects at the Allen Institute for Neural Dynamics for over a year. It has been used to sort data from over 1,000 sessions, finding over 1 million putatively neural units. This operational experience has helped refine the pipeline’s performance, robustness, and usability, positioning it as a mature and reliable tool for other research groups. The spike sorting pipeline is already being used by around 50 researchers at the Kempner Institute at Harvard University, and it is being adopted by other institutes as an end-to-end standardized spike sorting solution.

The results of the benchmarking pipeline Application #1 motivated our decision to replace `Kilosort2.5` with `Kilosort4` as the default spike sorter in our spike sorting pipeline. We found that `Kilosort4` could more accurately identify ground truth spikes (Figure 7), which corroborated the results of the original paper describing this algorithm (Pachitariu et al. 2024). We extended this result to data with the same noise characteristics and spiking statistics as what we were actively collecting, as well as to data from Neuropixels 2.0, which differs in its signal bandwidth and bit depth from Neuropixels 1.0.

We also used the benchmarking pipeline to test the impact of lossy compression on spike sorting accuracy (using `Kilosort4`). We previously showed that the `WavPack` audio compression algorithm, operating in lossless mode, achieved higher compression ratios for electrophysiology data than any general-purpose codecs (Buccino et al. 2023). In lossy mode, `WavPack` could reduce file sizes by more than 2x the lossless versions, with minimal distortion of spike waveform shapes. Such a decrease in file size has the potential to save many thousands of dollars in data storage costs, so there was a strong impetus to determine whether lossy compression has any adverse effects on downstream data processing. Here, we observe no impact on spike sorting performance after applying lossy compression to Neuropixels 1.0 data, and a slight degradation for Neuropixels 2.0 data. Even for Neuropixels 2.0, the overall impact on spike sorting quality is quite small, such that at the highest compression setting spikes are still detected more accurately by `Kilosort4` than by `Kilosort2.5` operating on lossless data (Figure 8a). The minimal impact of 7× compression ratios on sorter performance (Figure 8b) makes lossy compression with `WavPack` an attractive option as electrophysiology datasets expand to thousands of channels and across many days of continuous recording.

### Limitations of our approach

The majority of laboratories performing large-scale electrophysiology typically insert one or two probes per experiment. Therefore, when running spike sorting on local workstations with a single GPU, they will experience minimal speedup from using our pipeline in comparison to calling `SpikeInterface` directly. In addition, `SpikeInterface` is implemented in pure Python, whereas `Nextflow` depends on a Java installation (no longer available by default on most operating systems), and requires Windows Subsystem for Linux to run in a Windows environment. However, one option to bypass installation issues is to run the main pipeline script in container images pre-packaged with `Nextflow` (https://hub.docker.com/r/nextflow/nextflow). That said, for many scientists, the benefits of standardization and reproducibility offered by our pipeline will more than justify the additional setup and run time. The ability to trace the exact code and parameters used to generate a spike sorting result represents an important step toward transparent and publishable analyses. Moreover, access to a wide range of visualization tools and quality metrics enables rigorous data validation and facilitates meaningful comparisons across labs.

While we sought to provide a full end-to-end efficient solution, the presented pipelines might still seem overly complex and inflexible to some users. In part, this is due to the need to encapsulate each process in a separate GitHub repository pinned to a specific commit, which makes the actual executed code harder to find. We tried to address this limitation by providing extensive documentation on the pipeline architecture, which includes direct links to the individual repositories associated with each process. To add flexibility to the pipeline, we made sure key parameters are exposed and documented, and that these can be modified at runtime via an input JSON file or command-line arguments. Users can override the pipeline’s default preprocessing steps by providing a preprocessing pipeline dictionary as an input argument. It is also possible to limit the number of postprocessing extensions, for example by skipping expensive computations involved in computing spike locations or quality metrics. Downstream pipeline steps are resilient to these changes, e.g., the curation step will not attempt to run if quality metrics are not computed. Finally, the documentation includes instructions on interfacing with new spike sorting algorithms, with a GitHub template repository that can be used as a starting point. Currently, adding a new sorter is not entirely straightforward for end users. As a result, the core maintainers plan to add support for new sorters as they become available. All pipeline customization options can be found in the pipeline documentation: https://aind-ephys-pipeline.readthedocs.io/en/stable/customization.html.

Another limitation is related to integration with tools for manually inspecting spike sorting outputs. Currently, the spike sorting pipeline creates `sortingview` links that can be used to remotely view and curate spike sorting outputs. However, many users may prefer to perform curation on a local workstation with tools such as `phy` (Rossant et al. 2025) or `JRClust GUI` (Jun et al. 2025), which are fast and provide access to the raw data. While `SpikeInterface` does provide the ability to export data into `phy` format, our pipeline does not natively support `phy` or other desktop visualization tools. To mitigate this issue, we are taking a two-pronged approach. First, `SpikeInterface` should integrate the latest tools for automated curation, such as cluster labeling and merging/splitting. In many cases, existing tools such as `UnitRefine` already do most of the labeling that was previously done manually. LLM-based tools could also provide guidance for automated curation (Lin et al. 2025). Secondly, the pipeline is fully compatible with the `SpikeInterface-GUI` (Garcia et al. 2026), a desktop- and web-based curation interface with full access to raw data, which would further enhance usability and encourage broader adoption. `SpikeInterface-GUI` can readily open the postprocessed data generated by the spike sorting pipeline, and already includes many of the features of the previously mentioned tools.

The hybrid evaluation approach also presents some limitations. First, the ground truth units had Poisson-distributed spike times, which do not necessarily match the overall firing statistics of the original recording. One improvement to this approach could be to estimate ongoing population firing rates and inject spike trains that follow the dynamics of nearby neurons. The placement of hybrid units could also be chosen to match the observed neuronal density of the underlying recording, rather than using a uniform distribution. A second limitation lies in the selection of the spike templates: although we used different templates for each probe type, we did not choose templates to match the brain region of the original recording. It remains to be tested whether generating more realistic hybrid recordings will have any effect on spike sorting accuracy.

### Future directions

Looking ahead, we plan to expand our spike sorting pipeline’s capabilities with advanced curation workflows, broader support for emerging processing and spike sorting methods, and streamlined deployment across diverse backends. As growing datasets make manual curation increasingly impractical, such automated and standardized evaluation will become even more essential. The benchmarking pipeline will continue to develop as an open evaluation framework, enabling transparent and reproducible comparisons of spike sorting and preprocessing methods across the community. As one example, the work on lossy compression could be extended with additional codecs and parameter settings, exploiting our ability to read out spike sorting degradation directly from the hybrid ground truth spike times.

Although we ultimately view full automation of spike sorting as necessary for reproducibility, the field is not yet at that point. For this reason, we include interfaces for manual curation as part of our core pipeline. We anticipate that automated approaches will soon eliminate the need for human intervention, and the flexibility of our framework will allow these methods (as well as new spike sorting algorithms) to be readily incorporated. While the pipeline currently only supports `UnitRefine`, we plan to add other automated tools for curation, such as `Bombcell` (Fabre et al. 2023) and `SLAy` (Koukuntla et al. 2025). Importantly, our benchmarking tools can be used to evaluate and compare the accuracy of different curation strategies that operate without a human in the loop.

By providing an open, transparent, and well-documented framework, we aim to advance reproducibility and efficiency in the analysis of large-scale electrophysiology data. We hope our pipelines will accelerate community-driven improvements in spike sorting algorithms and lower technical barriers to adopting the most up-to-date approaches. As the field moves toward even more comprehensive and integrative views of brain function, a robust spike sorting ecosystem is just as critical to progress as scaling up the number of channels on each recording device.

## Methods

### Spike sorting pipeline steps

#### Job Dispatch

The job dispatch step scans an input directory containing raw electrophysiology data from one session and creates a list of *job* JSON files that control parallelization. Each JSON file contains a `recording_dict`, a dictionary representation of the `SpikeInterface Recording` object. Each `Recording` represents data from one contiguously sampled interval and one device, also known as a “stream.” The JSON file allows downstream processes to easily reload the `Recording` (using the `spikeinterface.load()` function) while keeping the size of data passed through different processes minimal. For probes that acquire separate high-frequency (AP) and low-frequency (LFP) streams (e.g., Neuropixels 1.0), a link to both data files is stored in the JSON file, as this information is used at the NWB Packaging step. The JSON file also contains a `session_name` and `recording_name` that are used throughout the pipeline to track the session and recording identity.

The `input` parameter allows the user to specify one of five formats for the input data:

1. `spikeglx`
2. `openephys`
3. `nwb`
4. `aind`^4^
5. `spikeinterface`

For the `spikeinterface` input, the `spikeinterface_info` field provides further information on how to read the data and attach the required probe information. For example, here is a sample `spikeinterface_info` for a dataset recorded by a data acquisition system from Intan Technologies:

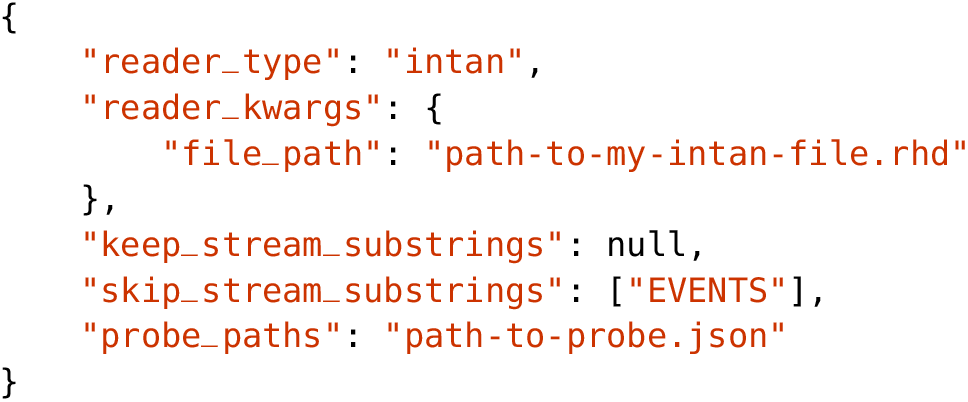

As a subset of stream may not contain neural data, the `keep/skip_stream_substrings` fields enable users to filter which streams to process with the full spike sorting pipeline (in the example above, streams with names containing the string “EVENTS” would be ignored). The `probe_paths` field allows users to specify paths to JSON files represented in `ProbeInterface` format to attach probe information to each recording (Garcia et al. 2022).

The input data folder can contain multiple *blocks* and *segments*, as defined in the `Neo`^5^ data model (Garcia et al. 2014). These concepts are inherited in `SpikeInterface`. A block typically refers to data from a single session or experiment (i.e., from one subject on one day). A segment represents data recorded contiguously within a block. For example, an `Open Ephys` input folder could contain multiple “experiments” (blocks) each with multiple “recordings” (segments)^6^ or an `NWB` file could contain several electrical series in its acquisition field. If the input data contains multiple blocks, each block will always be processed separately. For blocks with multiple segments, the user can choose whether to concatenate them (one JSON file is produced for the concatenated segments) or keep them separate (one JSON file is produced for each segment). The `split_segments` parameter controls this behavior.

In addition to producing multiple JSON files for each block (and optionally segment), this step also provides the option to split the channels of a recording into groups (e.g., individual shanks) that will be sorted independently. Channel groups are loaded automatically in the case of Neuropixels input data (recorded with `Open Ephys` or `SpikeGLX`) and also for `NWB` files (where this information is stored in the `electrodes` table). For input data loaded in `spikeinterface` format, the channel groups can be specified in the probe JSON file.

In addition, the job JSON files also include a *debug* flag, to communicate that the pipeline was started in debug mode, and a *skip times* flag that is set to true if the timestamps are corrupted and non-monotonically increasing, which is used to inform the NWB packaging step to reset the timestamp information.

The GitHub repository associated with this step can be found at:

https://github.com/AllenNeuralDynamics/aind-ephys-job-dispatch

#### Preprocessing

The preprocessing step is the first step that runs in parallel. A separate preprocessing instance is spawned for each JSON file produced by the job dispatch step. First, the preprocessing step reloads the `Recording` object from the JSON file. The recording then undergoes four transformations: phase shift, filtering, denoising, and motion estimation/correction.

The phase_shift function was originally developed by the International Brain Laboratory (IBL) (International Brain Laboratory 2024). The correction for these channel-wise “delays” is performed in the frequency domain, where a time delay corresponds to a phase shift rotation. The signal is therefore first FFT-transformed, then the delay is applied by rotating the phase of the signal in the frequency domain by 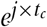 where *t*_*c*_ is the inter sample shift for channel *c*, which represents the relative delay within the sampling period (between 0 and 1). Finally the FFT transform is inverted to return to the time domain. For Neuropixels data, the *inter sample shift* property is loaded automatically from the probe metadata. For other probe types, this property must be explicitly specified. Note that this step is equivalent to the `Tshift` step in `CatGT`, from the `SpikeGLX` tool (Karsh, n.d.).

For the filtering step, a 5th order Butterworth highpass_filter with cutoff frequency at 300 Hz is applied by default. Users can optionally select bandpass_filter with cutoffs at 300 Hz and 6 kHz. The filter type, cutoff frequencies, and filter orders can be modified by the user. Internally, the `SpikeInterface` functions use the `scipy.filtfilt` function that applies a forward and a backward pass to maintain zero phase.

The denoising step performs channel masking followed by global denoising. The detect_bad_ channels function is run using the `coherence+psd` method, which is ported from IBL (International Brain Laboratory 2024). “Dead” channels are detected based on a low similarity with surrounding neighbor channels. “Noise” channels are those whose power between 80% and 100% of the Nyquist frequency is above a certain threshold (default = 0.02 µV^2^ / Hz) and have a high coherence (>1) with nearby channels (normal channels have coherence around 0, see (International Brain Laboratory 2024)). “Out” channels are channels at the top of the probe with low similarity to other non-neighboring channels, and are assumed to be outside the brain. The final channel labels are obtained as a majority vote on labels from 100 300-ms chunks randomly sampled throughout the recording. All three types of channels (*dead, noise, out*) are removed from the recording by default. Optionally, the user can choose to keep *out* channels (`remove_out_channels` parameter) or to keep *dead*/*noise* channels (`remove_out_channels` parameter). If more than 50% of channels are removed from a given stream, further processing (including spike sorting) is skipped. Downstream processing is also skipped if the recording duration is too small (by default < 120 seconds). Both the fraction of remaining channels required to skip further processing and the minimum recording duration can be adjusted by the user. After removing channels, a global denoising step is performed via Common Median Reference (CMR) or highpass spatial filtering, also referred to as destriping (International Brain Laboratory 2024).

Finally, motion is estimated and optionally corrected for. The motion correction step uses the compute_motion function, which can estimate the motion from the recording using several presets. By default, the `dredge_fast` preset is used, which uses the `grid_convolution` method to speed up spike depth estimation and `DREDge` for motion estimation (Windolf et al. 2025). The `grid_convolution` method for spike localization, introduced by (Pachitariu et al. 2024), uses an exhaustive catalog of artificial templates at known positions to estimate the position of a given spike as the weighted average of the spike waveform projected onto the templates in this catalog. For more details on spike localization and motion correction methods in general, we refer the reader to (Garcia et al. 2024). By default, motion is only estimated and not “applied” to the traces. This is done to let the spike sorter perform its own motion correction. However, the user can opt to also apply motion correction (setting `preprocessing.motion` parameter to `apply`) or to skip this step entirely (setting `preprocessing.motion` parameter to `skip`). When motion is applied, the traces are interpolated in space using *kriging* interpolation depending on the motion signals (Garcia et al. 2024).

After preprocessing, the recording is saved to a temporary binary file to speed up downstream steps. This is done in parallel using 1-second chunks. To minimize artifacts at boundaries, especially due to filtering, each chunk is padded with 5 ms from adjacent chunks. Alongside the binary file, this step also produces JSON files that represent the *lazy* preprocessing chain applied before and after motion correction, to enable `SpikeInterface` to reload the preprocessed recording from the raw data.

The GitHub repository associated with this step can be found at:

https://github.com/AllenNeuralDynamics/aind-ephys-preprocessing

#### Spike sorting

The spike sorting step first reloads the preprocessed recording saved to the temporary binary file. If the recording is multi-segment and the spike sorter doesn’t support multi-segment recordings, the segments are concatenated. The spike sorter (loaded from a versioned `DockerHub` container image) is then run on the input data. After spike sorting is complete, the output is re-split into multiple segments and any units with no spikes are removed. Spikes exceeding the number of samples of the recording, if any, are also removed (with the remove_excess_spikes function). The spike sorting output (a [Sorting](https://spikeinterface.readthedocs.io/en/stable/api.html#spikeinterface.core.BaseSorting) object) is saved to a `SpikeInterface` folder with the `sorting.save(folder=“…”)` function, so that it can be easily reloaded.

There is a separate GitHub repository for the spike sorting step associated with each sorter currently supported by the pipeline:

- Kilosort4: https://github.com/AllenNeuralDynamics/aind-ephys-spikesort-kilosort4
- SpyKING CIRCUS2: https://github.com/AllenNeuralDynamics/aind-ephys-spikesort-spykingcircus2
- Kilosort2.5: https://github.com/AllenNeuralDynamics/aind-ephys-spikesort-kilosort25

#### Postprocessing

The postprocessing step takes the preprocessed recording and the spike sorting output as inputs. These are combined into a [SortingAnalyzer](https://spikeinterface.readthedocs.io/en/stable/api.html#spikeinterface.core.SortingAnalyzer), which is the central object for performing additional computations. At instantiation, the *sparsity* of each unit, i.e., the set of channels the unit is defined on, is calculated via a fast template estimation using 100 uniformly sampled spikes per unit. The sparsity includes the channels within a 100 µm radius from the channel with the largest amplitude (*extremum* channel). A first `SortingAnalyzer` object is created and used only to remove duplicated units (remove_duplicated_units function), which are pairs of units that share over 90% of their spikes. We retain the unit whose spike times are best aligned to the template peak (knowing the cutouts used to compute the template). After duplicated units are removed, a new `SortingAnalyzer` runs the following *extensions*:

##### 1. random_spikes

Uniformly samples a certain number of spike for all units (default = 500).

##### 2. noise_levels

Randomly samples snippets of the preprocessed recording (default = 20 snippets of 500 ms each) and computes the median absolute deviation as a proxy of noise level for each channel.

##### 3. waveforms

Retrieves and saves sparse waveforms for the randomly sampled spikes (scaled to µV) and saves them as a 3D array with dimensions (`n_spikes`, `n_samples`, `n_max_sparse_channels`). The default cutout is -3 ms/+4 ms (i.e, 140 samples for a 30 kHz recording). Note that it might happen that some units, especially at the border of the probe, have fewer sparse channels than others: in this case, the waveforms array is padded with `NaN` values.

##### 4. templates

Computes each unit’s template as the mean of the sampled waveforms (when `waveforms` are available). In addition, it also computes the median and standard deviation of the waveforms.

##### 5. template_similarity

Computes the pairwise template similarity using L1 distance. This metric is preferred over other similarity metrics, such as cosine similarity, because the latter is not sensitive to amplitude differences.

##### 6. correlograms

Computes auto- and cross-correlograms for each pair of units using a bin size of 1 ms and a window of ±50 ms.

##### 7. isi_histograms

Computes inter-spike-interval histograms for each unit using a bin size of 5 ms and a window of 100 ms.

##### 8. unit_locations

Uses the unit templates to estimate the x, y, and z location with respect to the probe. This is done using the monopolar triangulation method, which models the peak-to-peak amplitudes of the templates on each channel as being generated by a current monopole:

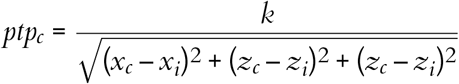

where *ptp*_*c*_ is the peak-to-peak amplitude of channel *c*; *x*_*c*_, *y*_*c*_, and *z*_*c*_ are the x, y, and z position of channel *c, x*_*i*_, *y*_*i*_, and *z*_*i*_ are the x, y, and z position of the unit *i*; and *k* is a term that includes the magnitude of the current and propagation properties of the tissue (see (Buccino et al. 2018) and (Garcia et al. 2024) for more details). The method finds the *k, x*_*i*_, *y*_*i*_, and *z*_*i*_ that minimize the error between the observed and reconstructed *ptp*_*c*_ on the sparse channels.

##### 9. spike_amplitudes

Retrieves the voltage value for each spike, sampling the traces at the extremum channel for each unit.

##### 10. spike_locations

Calculates a location for each spike using the grid convolution method, as explained in the **Preprocessing** section.

##### 11. principal_components

Uses the randomly sampled spikes for each unit to fit PCA (principal component analysis) models for each channel. This is done by incrementally fitting the waveforms from each unit on the channels they are defined on. After fitting the models, each of the randomly sampled waveform is projected on its sparse channels, so that the output is a 3D array with dimensions (`n_spikes`, `n_components`, `n_max_sparse_channels`).

##### 12. template_metrics

Computes several features that characterizes the templates. Prior to the computation, templates are upsampled by a factor of 10.

These metrics are computed from the extremum channel:

- `peak_to_valley`: spike duration between negative and positive peaks
- `halfwidth`: spike duration at 50% of the amplitude during the negative peak
- `peak_to_trough_ratio`: ratio between negative and positive peaks
- `recovery_slope`: spike slope between the negative peak and the next zero-crossing
- `repolarization_slope`: spike slope between the positive peak and the next zero-crossing
- `num_positive_peaks`: the number of positive peaks in the template
- `num_positive_peaks`: the number of negative peaks in the template

All durations are in seconds and slopes in µV / s.

In addition to these one-dimensional metrics (that use the template on a single channel), this extension also computes the following multi-channel metrics:

- `velocity_above`: the propagation velocity above the extremum channel of the template (over the probe depth direction)
- `velocity_below`: the propagation velocity below the extremum channel of the template (over the probe depth direction)
- `exp_decay`: the exponential decay of the template amplitudes over distance from the extremum channel
- `spread`: the spread of the template, computed as the distance of the Gaussian-filtered peak-to-peak amplitudes over depth (default smooth factor: 20 µm) which are above 20% of the maximum peak-to-peak amplitude.

Velocities are in µm / s, and spread and exponential decay are in µm.

For more information on template metrics, we refer the reader to the [SpikeInterface documentation](https://spikeinterface.readthedocs.io/en/stable/modules/postprocessing.html#template-metrics).

##### 13. quality_metrics

this extension computes the following quality metrics for each unit: *num spikes, firing rate, presence ratio, SNR, ISI violation, RP violation, sliding RP violation, amplitude cutoff, amplitude median, amplitude CV, synchrony, firing range, drift, isolation distance, l-ratio, d-prime, nearest neighbor, silhouette*.

For a detailed description of each quality metric, we refer the reader to the [SpikeInterface documentation](https://spikeinterface.readthedocs.io/en/latest/modules/qualitymetrics.html).

All parameters that control the computation of any extensions can be modified via the input parameter file. After all extension computations, the `SortingAnalyzer` object is saved in `Zarr` format.

The GitHub repository associated with this step can be found at:

https://github.com/AllenNeuralDynamics/aind-ephys-postprocessing.

#### Curation

The curation step uses the postprocessed data to apply labels to each unit. There are two set of labels, one based on quality metrics filters (`default_qc`), and another based on pre-trained classifiers (`decoder_label`). For `default_qc`, the user can input a `pandas.query` and use any combination of quality metrics computed at the postprocessing steps. The default query is:

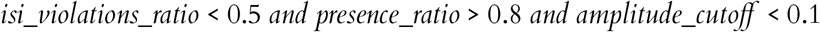

Units that satisfy this condition will have the `default_qc` value set to `True`.

The second set of labels come from `UnitRefine` (Jain et al. 2025), a set of pre-trained classifiers for labeling units after spike sorting. The classifiers were trained on hand-curated labels using quality and template metrics as input. The first classification is between *noise* and *neural* ([SpikeInterface/U-nitRefine_noise_neural_classifier](https://huggingface.co/SpikeInterface/UnitRefine_noise_neural_classifier)). For the units labeled as *neural*, a second classifier labels them as single versus multi-unit activity (SUA/MUA) (SpikeInterface/UnitRefine_sua_mua_classifier). `UnitRefine` labels are applied to the units using the auto_label_units function.

The GitHub repository associated with this step can be found at:

https://github.com/AllenNeuralDynamics/aind-ephys-curation.

#### Visualization

The visualization step uses `Figurl` and `sortingview` to produce shareable links that can be accessed by any device connected to the internet. This technology, integrated into `SpikeInterface`, pushes data needed for visualization to an accessible cloud location, whose Uniform Resource Identifier (URI) is embedded in the link itself. When following the generated links, the visualization plugin uses the URI to fetch the data from the cloud and provides rich and interactive tools to browse them. A simple installation step is required to set up this web visualization capability. We recommend creating a custom “zone” (i.e., providing one’s own cloud resources) if heavy usage is expected. More information can be found on the [Figurl documentation page](https://github.com/flatironinstitute/figurl/blob/main/README.md).

The visualization step generates two visualization links for each stream. The first link contains snippets of raw traces at different preprocessing stages, a raster plot of the complete session, and a visualization of the estimated motion signals. The second link includes two tabs: one with a web GUI for spike sorting visualization and curation, the second for exploring quality metric distribution and values for each units. Screenshots and links for example visualizations are available in Figure 3.

The GitHub repository associated with this step can be found at:

https://github.com/AllenNeuralDynamics/aind-ephys-visualization.

#### Quality control

The quality control step parallelizes over streams from the output of the Result Collection. For each stream, it produces the following metrics and figures:

##### Raw data

Plots 0.1 s AP and LFP traces at three points uniformly distributed over the recording duration. In case of wide-band input data, a 300 Hz highpass filter is used to extract the AP band, and a 0.1-500 Hz bandpass filter for the LFP band. These plots are used to inspect raw traces for artifacts, noise, and spiking activity.

##### PSD

Displays the Power Spectral Density of all channels both as a line plot and as a map, with the power over channel depth. The PSD is computed at three points uniformly distributed over the recording duration from 1 s snippets of traces. Three figures are generated for each stream: the first uses the unfiltered raw signal, the second focuses on the low-frequency (showing the 0.1-100 Hz band after applying a 0.1-500 Hz bandpass filter), and the third focuses on high frequency (showing the 5-15 kHz band after applying a 3 kHz highpass filter). These plots can be used to identify localized noise sources, such as 50 or 60 Hz line contamination.

##### Noise

Computes the root mean squared (RMS) values of each channel from the raw data and preprocessed data (using 20 chunks of 500 ms randomly sampled across the recording). The RMS values are plotted as lines over depths and can be useful for spotting abnormal channels, confirming the presence of channels outside of the brain, and checking that denoising resulted in lower noise levels than the raw data.

##### Drift

Uses the preprocessed data and pre-computed motion estimation to plot a raster map with overlaid estimated motion signals. This can be used to assess the overall drift in the recording and whether motion correction methods accurately estimated it. (Figure 3c).

##### Saturation

Detects events in the raw data where the signal reaches to the maximum or minimum voltage that the acquisition system can record. It produces a raster plot of positive and negative saturation events over time and a count of such events. These plots are used to assess the presence and extent of artifacts that result in ADC saturation, assuming the threshold voltage is known.

##### Unit Yield

Summarizes the output of the spike sorting, postprocessing, and curation steps by plotting histograms of selected quality metrics and scatter plots of unit amplitudes and firing rates over depth. It also displays a summary of the number of units in each category detected by curation step (# noise, # SUA, # MUA, and # units passing default QC) (Figure 3e). These plots are used to assess the overall yield and quality of the spike sorting.

##### Firing Rate

Produces a plot with the population firing rate (all units pooled together) over time, which is used to assess whether there are intervals with abnormally high or low activity during the recording (e.g., due to seizure-like events).

All figures are saved as `.png` files in the `quality_control` folder in the results and organized into a `quality_control.json` file following the AIND Data Schema [QualityControl](https://aind-data-schema.readthedocs.io/en/stable/quality_control.html) specification (see **AIND Data Schema**).

The quality control step includes two processes: one for computing metrics and plotting (run in parallel for each stream), and a second one to collect and organize the QC for different streams.

The GitHub repositories associated with this step can be found at:

- https://github.com/AllenNeuralDynamics/aind-ephys-processing-qc
- https://github.com/AllenNeuralDynamics/aind-ephys-qc-collector

#### Result collection

The result collection step gathers the output from all streams and re-organizes it into the folders described in **Spike sorting pipeline output**. Importantly, this step propagates any labels computed by the curation step to the `SortingAnalyzer Zarr` folders, so that these objects include all the information and computation outputs from the pipeline.

The GitHub repository associated with this step can be found at:

https://github.com/AllenNeuralDynamics/aind-ephys-results-collector.

#### NWB Packaging

The `SpikeInterface` objects generated throughout the pipeline are converted to NWB using functions from `neuroconv` (Mayorquin et al. 2025). Each NWB file includes the following data:

- Device/Electrodes/Electrode groups: Relevant probe metadata is automatically fetched from the input data, including probe serial number, manufacturer, name, electrode locations, and channel groups (e.g., for multi-shank probes).
- LFP: By default, LFP traces are downsampled in space and time.
- Units: All units from different probes are combined into one table that stores spike times, waveforms, and quality/template metrics.
- Raw traces (*optional*): Raw traces are not stored by default, as doing so would greatly increase the size of the NWB file.

When the pipeline input data is not already in `NWB` format, it is recommended to add `subject.json` and `data_description.json` files following the AIND Data Schema specification^7^ to the input data folder, so that this information can be used to instantiate NWB files with the correct metadata. In case these files are missing, NWB files with mock metadata will be created instead.

Both LFP and raw traces (if saved) are compressed using the `Zstandard` codec (Facebook 2025) implemented via Blosc (The Blosc Development Team 2025), which showed the best performance among natively supported codecs in `Zarr` (Buccino et al. 2023).

The NWB export step is carried out by two separate processes. The first one generates base NWB files with session and subject information and adds data from the raw *ecephys* input folder (Device/Electrodes/Electrode groups, LFP and optionally raw traces); the second one adds the post-spike sorting and curation data (Units table).

The GitHub repositories associated with these two processes are:

- https://github.com/AllenNeuralDynamics/aind-ecephys-nwb
- https://github.com/AllenNeuralDynamics/aind-units-nwb

### Spike sorting pipeline output

The spike sorting pipeline output is organized into a single “results” folder with following sub-directories:

- `preprocessed/`: JSON files with preprocessed data for each recording stream, alongside subdirectories containing motion estimation results.
- `spikesorted/`: Raw spike sorting outputs.
- `postprocessed/`: `Zarr`-format `SortingAnalyzer` outputs with computed extensions.
- `curated/`: Information about unit deduplication and classification (e.g., noise, multi-unit activity, single-unit activity) labels. Properties such as `default_qc` and `decoder_label` are available for downstream analysis and validation.
- `visualization_output.json`: `Figurl` links for interactive visualization of each stream.
- `quality_control/ and quality_control.json`: QC visualizations and a JSON file following the `QualityControl` schema from the AIND Data Schema (see **AIND Data Schema**).
- `nwb/`: Neurodata Without Borders (NWB) files, with one file per block or segment. Each NWB file includes curated spike times and unit IDs, electrode metadata, and (optionally) local field potential (LFP) signals.
- `nextflow/`: Workflow metadata, provenance information, timeline, and execution logs.
- `processing.json`: Logs processing steps, parameters, and execution times, following the `Processing` schema from the AIND Data Schema specification (see **AIND Data Schema**).

### Evaluation pipeline steps

The evaluation pipeline re-uses the Job Dispatch and Spike Sorting (`Kilosort2.5` and `Kilosort4`) steps of the spike sorting pipeline. The three steps unique to the evaluation pipeline are described below:

#### Hybrid generation

The hybrid generation step begins by estimating motion of the input recording using `DREDge`, after light preprocessing (highpass and common median reference). Depending on the probe attached to the recording, which is part of the `Recording` metadata, the templates available through the `SpikeInterface` template library are fetched (with the fetch_templates_database_info() function) and pre-filtered by probe type.

For Neuropixels 1.0 recordings, templates from the International Brain Lab (IBL) Brain-Wide Map (International Brain Laboratory et al. 2024; International Brain Laboratory et al. 2025) are selected (2,183 templates from 37 different insertions). For other types of probes, including Neuropixels 2.0, high-density Neuropixels Ultra templates from (Steinmetz & Ye 2022) are used (5,694 templates).

Next, we randomly select *T* templates out of the pre-filtered templates by probe type (*T* = 10 by default). For probes other than Neuropixels 1.0, which use the Neuropixels Ultra templates, an additional step is added to interpolate the templates (using *kriging*) to the target probe geometry, and to randomly relocate them between the 5^*th*^ and 95^*th*^ percentiles of the distribution of detected peak depths (detected during the motion estimation process). This is to ensure that hybrid units and spikes fall within an active probe region. For Neuropixels 1.0, neither interpolation nor relocation is performed, since templates cover the entire probe span. In this case, the template selection is performed so that the selected templates are between the 5^*th*^ and 95^*th*^ percentiles of the peak depth distribution. Once templates are selected (and interpolated/relocated for Neuropixels Ultra), they are finally scaled to a user-defined range, which is [50, 200]µV by default. These steps are repeated such that *IT* random iterations of hybrid recordings (*IT* = 5 by default) are generated for the same input dataset.

The GitHub repository associated with this step can be found at:

https://github.com/AllenNeuralDynamics/aind-ephys-hybrid-generation

#### Hybrid evaluation

The hybrid evaluation step collects and reorganizes all generated hybrid data and spike sorting results from the different spike sorting cases. The output files for each session are organized to mimic a [spikeinterface.benchmarks.SorterStudy](https://github.com/SpikeInterface/spikeinterface/blob/main/src/spikeinterface/benchmark/benchmark_sorter.py#L45) - generated folder, in order to exploit all its comparison and plotting capabilities. In doing so, the spike sorting output of each spike sorting case is compared against its hybrid ground-truth. The `SorterStudy` object then generates several `pandas.DataFrames` for performance, run times, unit classification counts (well detected, redundant, overmerged, false positive; see (Buccino et al. 2020)) that are saved as CSV files and plotted. Finally, all CSV from all sessions are combined to generate aggregated plots using comparison data from all sessions (e.g, Figure 7, Figure 8a).

The GitHub repository associated with this step can be found at:

https://github.com/AllenNeuralDynamics/aind-ephys-hybrid-evaluation

#### Compression

For the lossy compression application, the spike sorting cases include lossy compression at various levels, preprocessing (described above), and spike sorting with `Kilosort4` (described above).

The compression step simply loads the recording and saves it to `Zarr` using the `SpikeInterface` compression framework. The `Zarr` compressor is `WavPack` from the `wavpack-numcodecs` package^8^, instantiated with different `bps` levels for each spike sorting case (3, 2.5, 2.25) (Buccino et al. 2023).

The GitHub repository associated with this step can be found at:

https://github.com/AllenNeuralDynamics/aind-ephys-compress

### Evaluation pipeline output

The evaluation pipeline output includes the following files and folders. All of the folders are organized per session, except for the `aggregated/` folder which aggregates all the results and plots from all input sessions.

- `figures/`: Visualizations of the various steps of the pipeline:
  – `benchmarks/`: Figures to display the performance of the different spike sorting cases, including overall performance curves (accuracy, precision, and recall) ordered by performance and by signal-to-noise ratio, run times, and unit classification counts.
  – `templates/`: Figures to show all selected and interpolated hybrid templates, to assess the quality of the hybrid units.
  – `rasters/`: Figures with raster maps of the original recording, with overlaid rasters of hybrid spikes and motion estimates.
  – `motion/`: Figures with summary motion signals.
- `dataframes/`: CSV files with unit-level performance (accuracy, precision, recall), unit counts (well-detected, false positive, redundant, etc.), pre-computed quality metrics, and run times
- `motion/`: Estimated motion data from `SpikeInterface`.
- `gt_studies/`: `SpikeInterface SorterStudy` folders, used to reload the comparison objects and perform further analysis.
- `aggregated/`: Aggregated dataframes and summary figures pulling all results from all sessions.
- `nextflow/`: Workflow metadata, provenance information, timeline, and execution logs.

### Statistical analysis

We used the non-parametric tests for all comparisons, given that all tested distributions were not normal. We used the Wilcoxon signed-rank test for paired samples (Figure 7a, Figure 8a) and Mann–Whitney U test for unpaired samples (Figure 7c-d, Figure 8c-d). For population tests (more than two samples), we used the non-parametric Kruskal–Wallis test. Post-hoc tests were run with Mann–Whitney U test for independent samples, and Wilcoxon signed-rank tests for paired samples with Holm’s correction for *P*-values.

Tests were performed using `scipy` (Virtanen et al. 2020) and `scikit-posthocs` (Terpilowski 2019). In addition to reporting *P*-values for two-sample tests, we also report the Cohen’s *d* coefficient as a proxy for the effect size.

#### Software versions

We list here the software versions used to run and generate the results. We refer the reader to the software documentation for updated supported versions.

`SpikeInterface` : v0.102.3

`Kilosort2.5` : v2.5.1

`Kilosort4` : v4.0.30

`WavPack` : v5.7.0

## Data and code availability

Hybrid data for evaluating spike sorters and lossy compression are available at the following URL: https://registry.opendata.aws/allen-nd-ephys-evaluation/

The code for the presented pipelines is available on GitHub:

- Spike sorting pipeline: https://github.com/AllenNeuralDynamics/aind-ephys-pipeline
- Evaluation pipeline: https://github.com/AllenNeuralDynamics/aind-ephys-hybrid-benchmark

The figures from the results section can have been generated with the following Code Ocean capsule: https://codeocean.allenneuraldynamics.org/capsule/9554286/tree/v4. The spike sorting pipeline is also available in the Code Ocean Public Collections: https://codeocean.allenneuraldynamics.org/capsule/4364160/tree/v4.

## Acknowledgments

We thank Heberto Mayorquin, Pierre Yger, Samuel Garcia, and Ben Dichter for the contributions to the development of the hybrid recording generation framework in `SpikeInterface`. We thank Zhiwen Ye, Nick Steinmetz, and the International Brain Laboratory for collecting and packaging the single unit templates. We thank Dan Birman and Carter Peene for their contributions to the AIND quality control portal. We thank Corbett Bennett, Hannah Cabasco, Jackie Kuyat, Vayle LaFehr, Ethan McBride, Ben Hardcastle, Sam Gale, Shawn Olsen, and Xinxin Yin for providing the raw data used to generate hybrid recordings. We thank members of the Allen Institute for Neural Dynamics for using the sorting pipeline to process their datasets and providing feedback on pipeline outputs and visualizations. We thank Bala Desinghu, Max Shad, Janet Wallace, and Bernardo Sabatini at the Kempner Institute at Harvard University for testing the sorting pipeline and providing statistics on their internal usage.

## Author contribution matrix

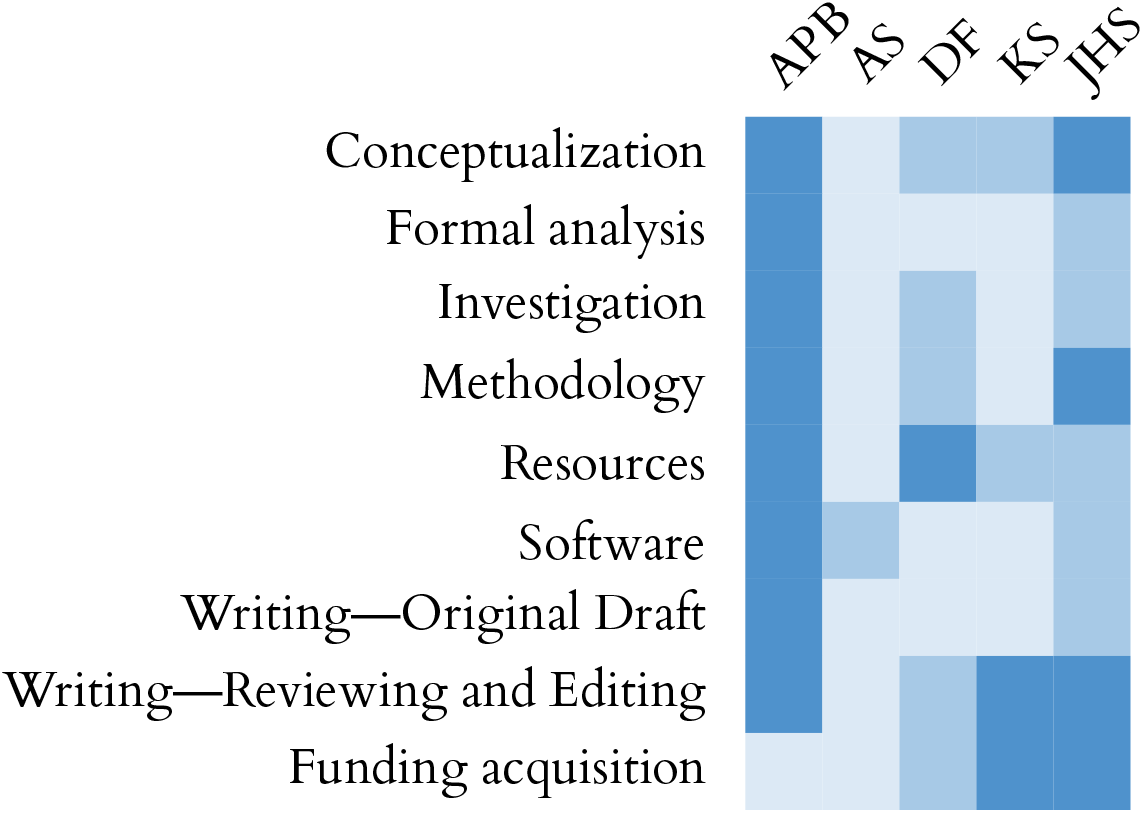

## Appendix 1. AIND Data Schema

The [AIND Data Schema](https://aind-data-schema.readthedocs.io/en/stable/) defines the specification for metadata that accompanies all data produced by the Allen Institute for Neural Dynamics. The goal of the schema is is to make all data consistently findable and understandable. The schema supports standard descriptions of the following categories of information:

- **Data Description**: administrative metadata about the source of the data, funding, relevant licenses, and restrictions on use.
- **Subject**: Species, genotype, age, sex, and source.
- **Procedure**s: descriptions of any procedures performed prior to data acquisition, including subject procedures (surgeries, behavior training, etc.) and specimen procedures (tissue preparation, staining, etc.).
- **Instrument**: descriptions of the equipment used to acquire data, including part names and serial numbers.
- **Acquisition**: how devices were configured during active data acquisition.
- **Processing**: descriptions of how data has been processed and analyzed into derived data assets, including information on the software and parameters used.
- **Quality Control**: Metrics and figures describing the quality of a data asset.
- **Model**: Metadata describing machine learning models created from or used to analyze data assets.

AIND Data Schema is written as a semantically versioned Python library. AIND Data Schema utilizes [Pydantic](https://docs.pydantic.dev/latest/), a common data validation library that integrates well with large variety of tools in the Python ecosystem. Pydantic is able to render AIND Data Schema to [JSONSchema](https://json-schema.org/) to support data validation outside of Python. Basic controlled vocabularies (list of organizations, instrument manufacturers, etc) are stored in a separate library called aind-data-schema-models.

For reference, a subject described in AIND Data Schema 2.0 looks like this:

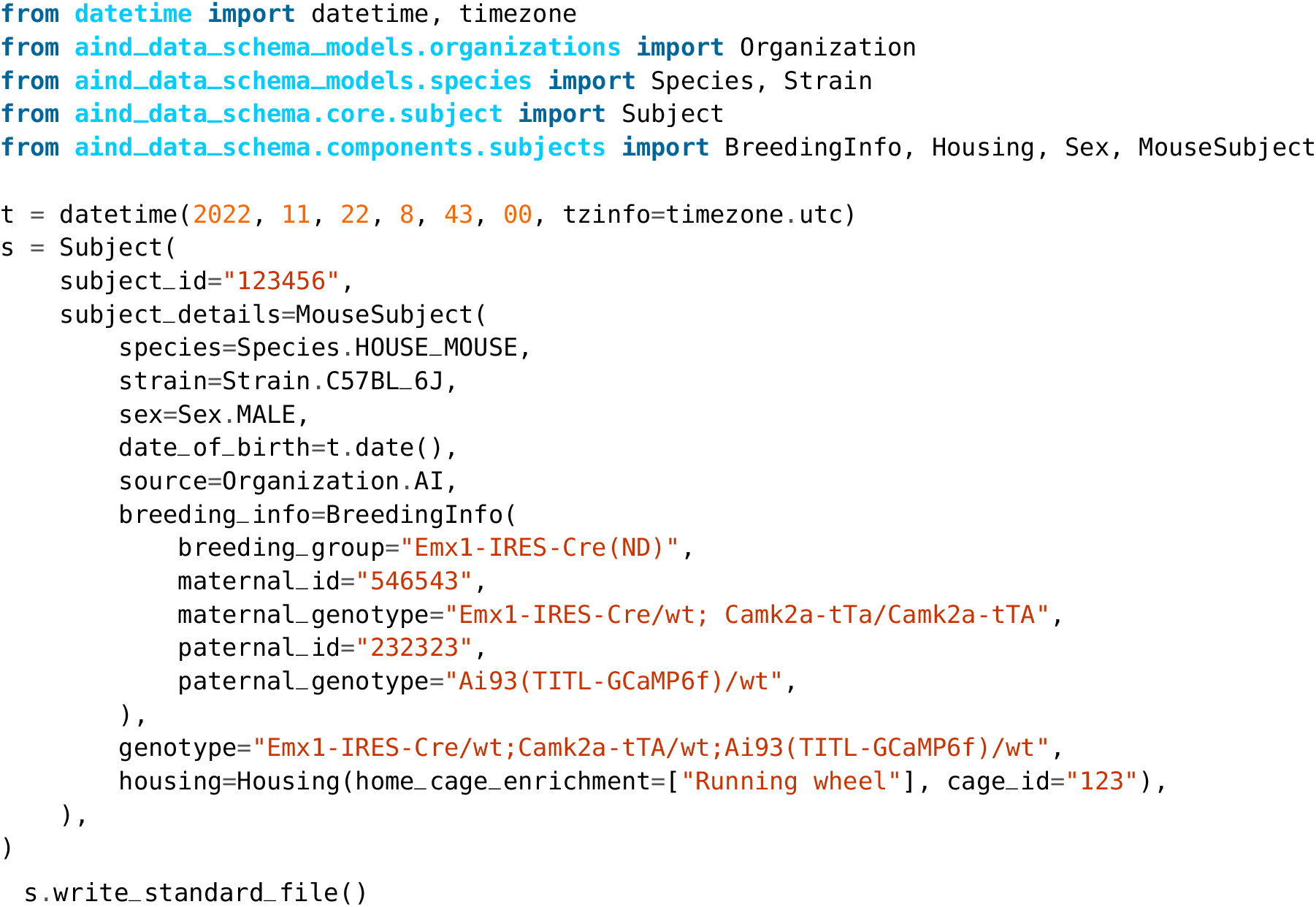

This produces `subject.json` in the script’s current working directory. Should there be validation errors (missing required information, improperly formatted data), AIND Data Schema will report these automatically.

---

**Figure S1:**
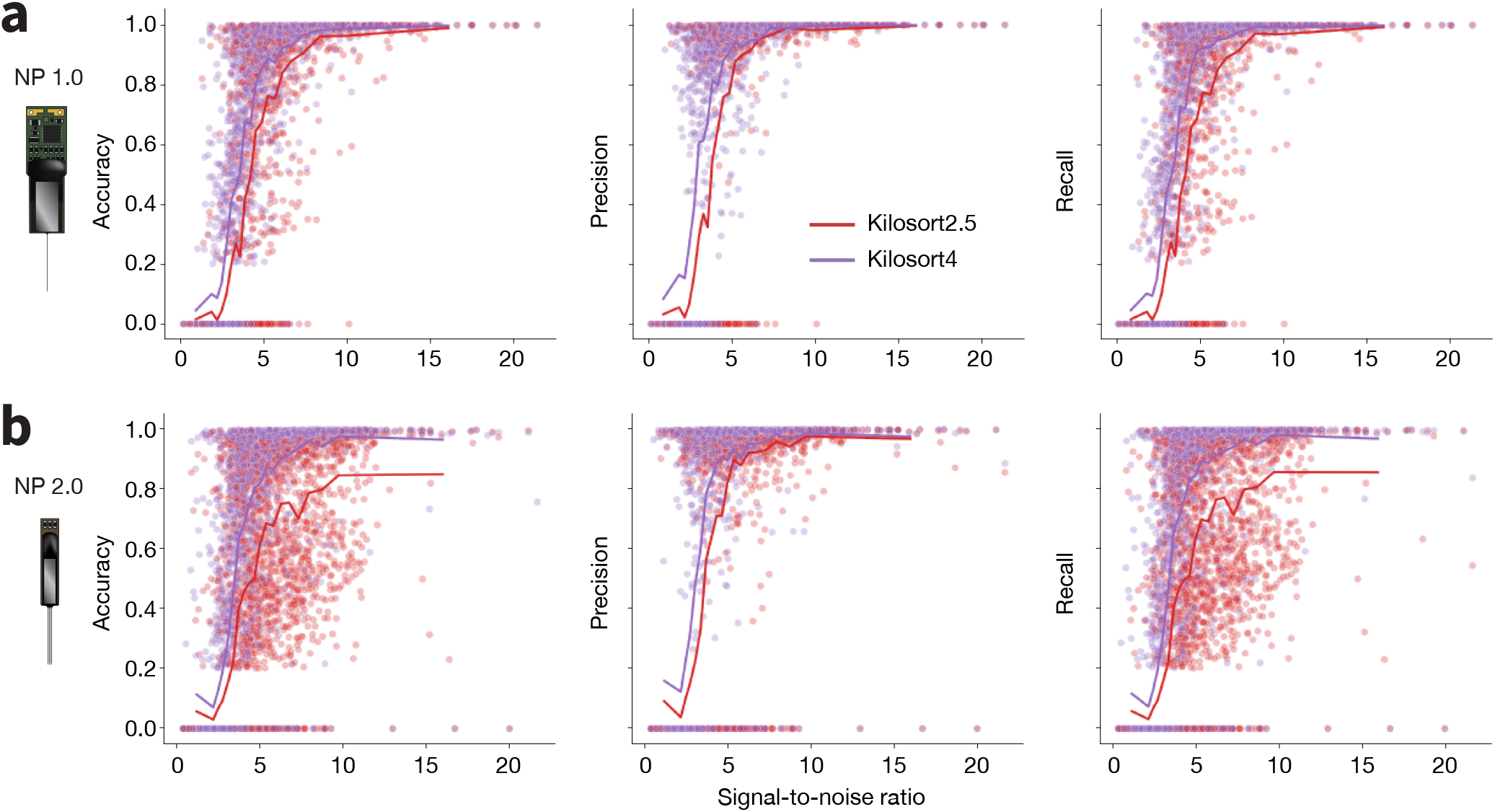
Spike sorter performance as a function of unit signal-to-noise ratio. **a**, Accuracy, precision, and recall for every hybrid units added to the Neuropixels 1.0. Lines indicate the average value at each signal-to-noise level. **b**, Same as a, but for Neuropixels 2.0 datasets.

**Figure S2:**
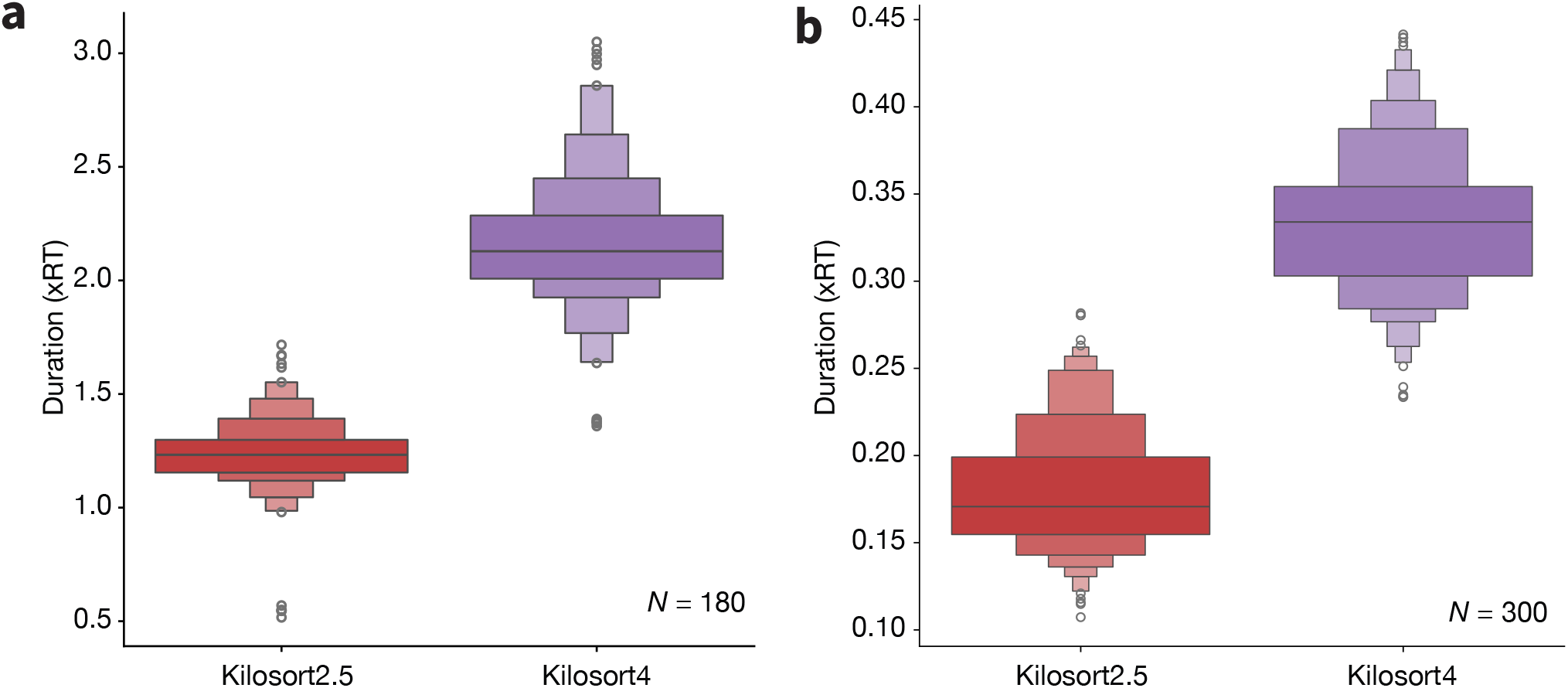
Kilosort run times. **a**, Run times of `Kilosort2.5` (red) and `Kilosort4` (purple) on Neuropixels 1.0 datasets. Run times are relative to real-time (xRT). b, Same as a, but for Neuropixels 2.0 datasets. Note that for Neuropixels 1.0, spike sorting was run on the full 384-channel recording, while it was split by shank (96 channels each) for Neuropixels 2.0, which is why the latter probe has faster execution times.

**Figure S3:**
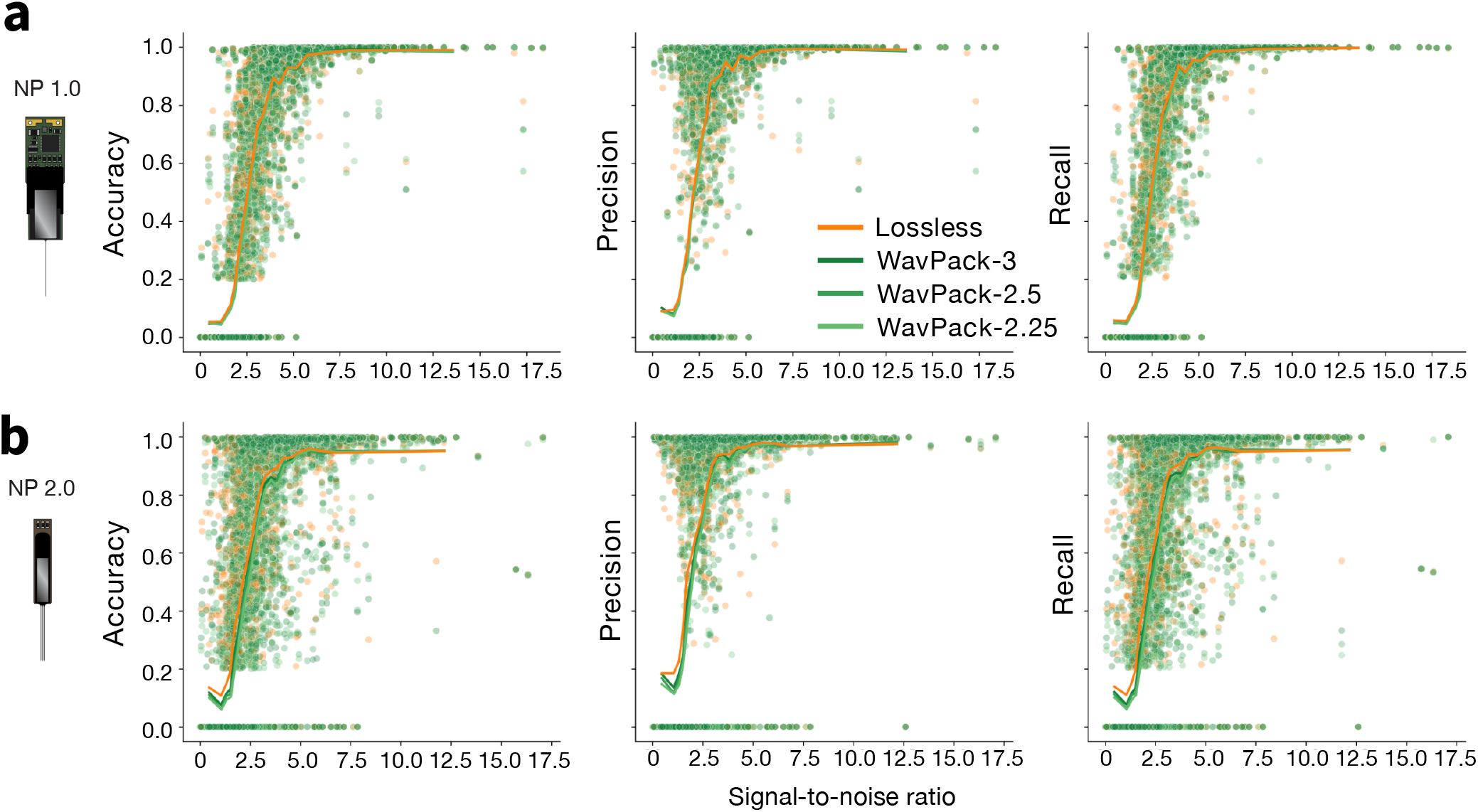
Lossy compression performance as a function of unit signal-to-noise ratio. **a**, Accuracy, precision, and recall for hybrid units added to Neuropixels 1.0 datasets. Lines indicate the average value at each signal-to-noise level. **b**, Same as a, but for Neuropixels 2.0 datasets.

## Footnotes

1 Total Run Time = [Non-Parallel] + [Parallel] = [(0.05 + 0.38 + 0.16) *×* 6 (probes) *×* 2 (hours)] + [(0.84 + 2-7 + 1. + 0.1 + 1.02) *×* 2 (hours)] = 18.40 hours

2 Total Run Time = 5.554 USD *×* 6 (probes) *×* 2 (hours) = 66.65 USD

3 The hybrid data for `Kilosort2.5` were generated with a different random seed, but with the exact same parameters.

4 `aind` is an internal format used at AIND which includes an `Open Ephys` folder with clipped binary files, `Zarr`-compressed traces, and metadata files

5 https://neo.readthedocs.io/en/stable/

6 A new recording is started whenever the experimenter pauses and then resumes the acquisition. A new experiment is made when the experimenter stops and restarts acquisition.

7 [Subject](https://aind-data-schema.readthedocs.io/en/stable/subject.html); [DataDescription](https://aind-data-schema.readthedocs.io/en/stable/data_description.html)

8 https://github.com/AllenNeuralDynamics/wavpack-numcodecs

